# EasyGuide plasmids support in vivo assembly of gRNAs for CRISPR/Cas9 applications in *Saccharomyces cerevisiae*

**DOI:** 10.1101/2022.07.01.498453

**Authors:** Ana P. Jacobus, Joneclei A. Barreto, Lucas S. de Bem, Yasmine A. Menegon, Ícaro Fier, João G.R. Bueno, Leandro V. dos Santos, Jeferson Gross

## Abstract

Most CRISPR/Cas9 applications in yeast rely on a plasmid-based expression of Cas9 and its guide RNA (gRNA) containing a 20-nucleotides (nts) spacer tailored to each genomic target. The lengthy assembly of this customized gRNA requires at least 3–5 days for its pre-cloning in *Escherichia coli*, purification, validation, and co-transformation with Cas9 into a yeast strain. Here, we constructed a series of 12 EasyGuide plasmids to simplify CRISPR/Cas9 applications in *Saccharomyces cerevisiae*. The new vectors provide templates for generating PCR fragments that can assemble up to six functional gRNAs directly into yeasts via homologous recombination between the 20-nts spacers. By dispensing pre-cloning in *E. coli*, yeast in vivo gRNA assembly significantly reduces the CRISPR/Cas9 experimental workload. A highly efficient yeast genome editing procedure, involving PCR amplification of gRNAs and donors, followed by their transformation into a Cas9-expressing strain, can be easily accomplished in a single day.

## INTRODUCTION

CRISPR/Cas9-mediated applications, such as genome editing, interference, and activation, belong to the modern *S. cerevisiae* genetics toolkit.^1^ Key components of the CRISPR/Cas9 system are the S*treptococcus pyogenes* Cas9 and a gRNA that directs the Cas9 into a specific genome location.^2^ This is possible through a 20-nts base pairing of the gRNA spacer with a sequence upstream to a NGG trinucleotide, the protospacer adjacent motif (PAM). Expression of native or mutated Cas9 proteins and gRNAs is usually based on shuttle plasmids that are first cloned in *E. coli* and then mobilized into *S. cerevisiae*.^3^ The gene specifying the Cas9 protein is encoded within most CRISPR/Cas9 vector backbones and is a standard component of the system. In contrast, the gRNA is a variable part of CRISPR/Cas9 plasmids and must be newly cloned for each experiment, as it contains the 20-nts spacer defining a particular Cas9 target.^3^ Therefore, a lengthy preparatory step is required, during which one or more gRNAs are usually ligated in vitro and transformed into *E. coli* for propagation.^3^ The proper assembly of gRNA(s) into the vector is usually authenticated by PCR or restriction analysis after DNA plasmid extraction. The whole procedure can take 3–5 days or even longer if Sanger sequencing is used to further confirm the correct plasmid construction. Only then is the gRNA-containing plasmid transformed into *S. cerevisiae*.

Several cloning methods and plasmid designs have been developed to simplify pre-cloning of single or multiple gRNAs; these range from traditional DNA cleavage and ligation^4-6^ to modern approaches such as USER cloning,^7^ Golden Gate assembly,^8-10^ or Gibson assembly.^11-13^ However, all these procedures generally require cloning kits, DNA modification enzymes, and/or special oligonucleotides that inevitably increase the costs of CRISPR/Cas9 experiments. An alternative tool is provided by *S. cerevisiae*, which can drive homologous recombination between DNA fragments whose overlaps are as short as 20 nts.^14^ In this way gRNA-containing plasmids can be directly assembled in vivo into the yeast targeted for CRISPR/Cas9 applications; eliminating the *E. coli* gRNA pre-cloning step reduces time and costs. Such advantages have inspired new protocols for homology-based gRNA assembly into yeasts.^11,15^ Though successful, most proposed methods are restricted to assembling single or duplex gRNAs^11,12,15,16^ or rely on preparatory steps, such as linearization of destination vectors,^13,15-17^ or gRNA pre-assembly by a fusion PCR.^18,17^ In vivo assembly of marker plasmids were also attempted in parallel to the expression of gRNAs from linear cassettes ordered from DNA synthesis companies.^19^ Despite these excellent approaches, a streamlined procedure based on standard plasmids to amplify PCR modules has not yet been developed to engineer multiple gRNAs in one step directly into the target yeast. Here, we introduce a versatile set of 12 plasmids (the EasyGuide series) to overcome these limitations, save costs, and reduce the CRISPR/Cas9 workflow. The proposed plasmids allow facile generation of PCR parts for assembly via yeast homologous recombination of up to six functional gRNAs.

## RESULTS AND DISCUSSION

### EasyGuide method overview

Our method is based on the Cas9 and gRNAs expression from different plasmids that are sequentially transformed (Figure 1A). The pEasyCas9 is a plasmid that specifies geneticin resistance and constitutively expresses the *S. pyogenes* Cas9 codon optimized for *S. cerevisiae*^11^ (Figures 1A and S1). The pEasyCas9 can be transformed only once and be maintained in a host strain repeatedly used for CRISPR/Cas9 applications (the Cas9 pre-loading mode). Alternatively, pEasyCas9 can be co-transformed with the donor(s) (i.e., the DNA repair template(s) for genome editing) into a strain in which the gRNAs-encoding plasmids have been previously assembled in vivo (the gRNA pre-assembling mode, Figure 1A). The pEasyG1, G2, and G3 are *E. coli* propagative plasmids that serve as templates to generate PCR fragments for assembling functional gRNAs in yeast (Figure 1B, 1C, and 1D). All PCR fragments are amplified from pEasyG1–3 plasmids with a single set of core primers: gA is a forward primer that binds to the gRNA scaffold, and gB is a reverse primer annealing to the SNR52 promoter (pSNR52) used for gRNA expression. The 20-nts spacer sequence for each planned Cas9 target is specified at the 5’ sequence of both gA and gB oligonucleotides, forming stretches of overlapping DNA that serve as substrates for in vivo homologous recombination (Figure 1B). The pEasyG-derived amplicons can be directly transformed into yeasts from the PCR reaction tubes without agarose gel or silica column purification and (except for pEasyG1 PCR reactions) dispensing DpnI treatment to eliminate the template plasmid.

**Figure 1.**
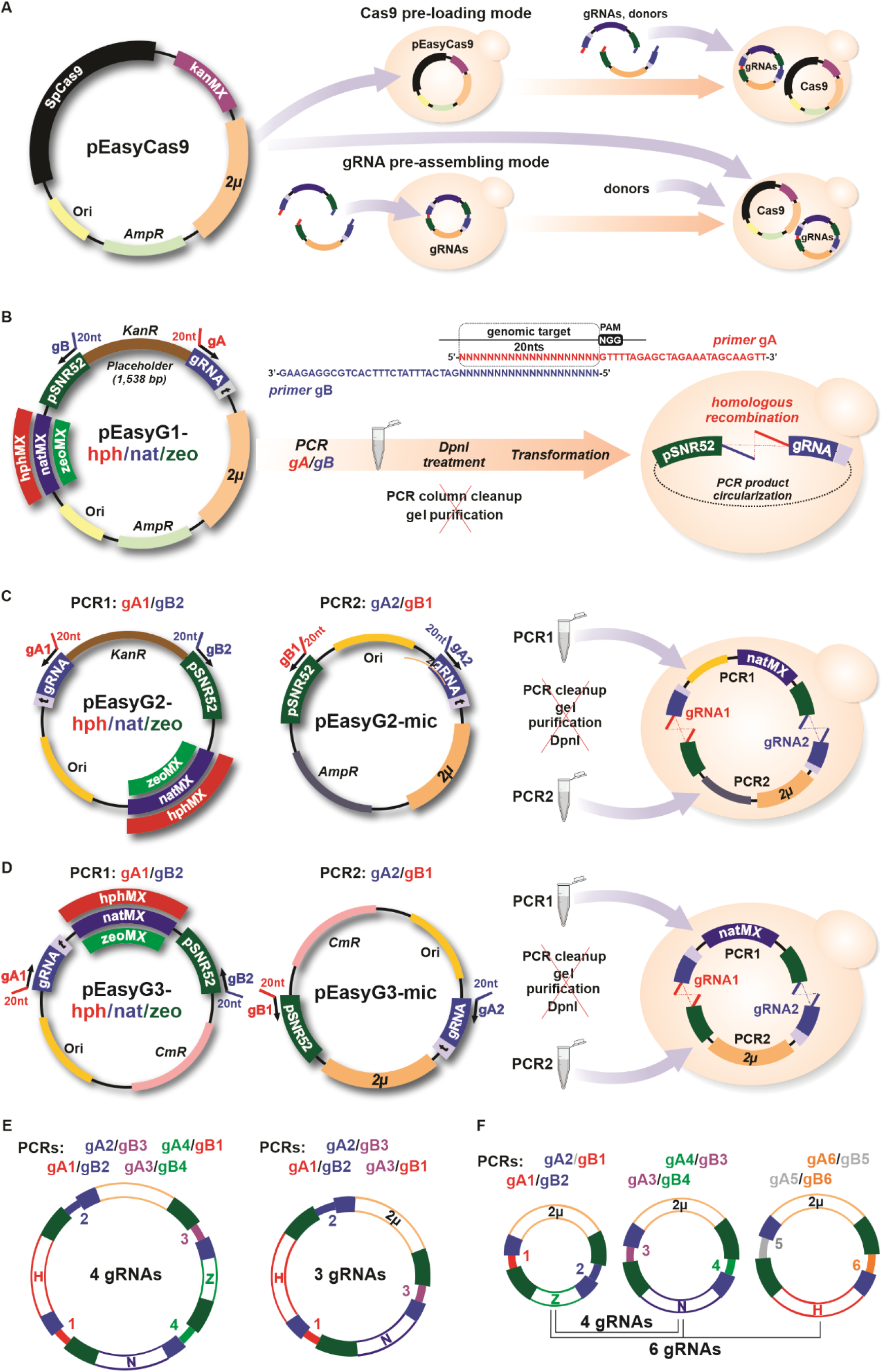
EasyGuide plasmids for yeast in vivo assembly of gRNAs. (A) The pEasyCas9 has a 2µ origin and encodes the *S. pyogenes* Cas9 and a geneticin resistance mark (kanMX). The pEasyCas9 can be inserted into a host strain that is used for genome editing experiments. Alternatively, a gRNA(s) may be pre-assembled into a yeast strain for subsequent co-transformation of pEasyCas9 and donor(s). (B) The pEasyG1 plasmids contain a gRNA scaffold (blue) and the Small Nucleolar RNA Pol III promoter (pSNR52, green) separated by a placeholder. pEasyG1 is a template to amplify PCR fragments with primers gA and gB. Both oligos specify 20-nts spacers that provide recombination sites for in vivo assembly of a functional gRNA. The spacer sequence of gA is taken from 20-nts upstream of the PAM sequence on the yeast genome. PCR fragments can be directly transformed from the PCR reaction into yeast cells, dispensing (red crossed) DNA purification steps. (C) The pEasyG2-zeo/nat/hph encode antibiotic resistance marks (zeoMX, natMX, or hphMX), and pEasyG2-mic contains the 2µ origin. In vivo recombination between amplicons (PCR1 and PCR2) derived from these templates generates a functional plasmid expressing two gRNAs. (D) Similarly, in vivo recombination between amplicons PCR1 and PCR2 from pEasyG3-zeo/nat/hph and pEasyG3-mic, respectively, results in a plasmid expressing two gRNAs. (E, F) Alternative ways to assemble three, four, or six gRNAs from pEasyG3(2) amplicons. PCR primer combinations are indicated. Plasmid features are not in scale.

### pEasyG1 plasmids

Each of the three pEasyG1 plasmids generates a PCR fragment that specifies a single gRNA (Figures 1B and S2). The pEasyG1-zeo/nat/hph encode antibiotic selection marks (zeocin, nourseothricin, and hygromycin, respectively) and the 2-micron (2µ) origin for propagation in *S. cerevisiae* (Figures 1B and S2). PCR fragments of 5,040 bp, 5,232 bp, and 5,688 bp are generated with gA/gB primers from pEasyG1-zeo, pEasyG1-nat, and pEasyG1-hph, respectively. The plasmid is circularized and the functional gRNA correctly assembled upon transformation of the PCR product into *S. cerevisiae* (Figure 1B). Up to three gRNAs can be simultaneously assembled into *S. cerevisiae* by co-transformation of PCR products generated from templates pEasyG1-zeo, -nat, -hph selected with their three respective antibiotics. The pEasyG1 template also bears a pUC19-derived ColE1 origin of replication and the ampicillin resistance marker^20^ (Figures 1B and S2) and therefore can also be circularized in *E. coli* by in vivo cloning^21^ or any chosen method. In this case, the placeholder (encoding a kanamycin resistance gene) provides a marker indicating false positives during bacterial cloning.

### pEasyG2 plasmids

This vector series provides templates to amplify complementary PCR fragments that recombine in vivo, yielding two functional gRNAs (Figures 1C, S3, and S4). PCR amplicons of 3,024 bp are generated from pEasyG2-mic with the gA/gB primer pair and contain at each extremity a 20-nts spacer. On one side the spacer is fused to the gRNA scaffold, whereas on the other end the spacer is connected to the pSNR52 (Figure 1C). Importantly, amplicons derived from pEasyG2-mic contain the 2µ sequence for replication in *S. cerevisiae* but lack a yeast selection marker. The PCR fragments from pEasyG2-zeo, -nat, -hph plasmids (2,453 bp, 2,645 bp, and 3,101 bp, respectively) are a counterpart antibiotic selection marker and lack the 2µ origin of replication. pEasyG2-derived PCR fragments also carry 20-nts spacers at their extremities fused with the gRNA scaffold and the pSNR52. Therefore, homologous recombination between the 20-nts spacers of PCR fragments (produced with primer pairs gA1/gB2 and gA2/gB1 from pEasyG2-mic and pEasyG2-zeo/nat/hph, respectively) generates a yeast replicative plasmid and two functional gRNAs (Figures 1C and S4B). Assembling plasmids from complementary essential modules is optimal because this method dramatically reduces cloning background and does not require treating PCR products with DpnI.^22^ Plasmid assembly in *E. coli* is a potential alternative because the pEasyG2-mic-derived PCR fragment carries an ampicillin selection marker, and amplicons from pEasyG2-zeo/nat/hph specify a pUC19-ColE1 origin of replication.

### pEasyG3 plasmids

Like the pEasyG2 series, amplicons generated from pEasyG3-mic (1,888 bp) recombine in vivo with pEasyG3-zeo/nat/hph PCR products (1,460 bp, 1,652 bp, 2,108 bp, respectively) due to the homology between 20-nts spacers specified by the gA1/gB2 and gA2/gB1 primer pairs (Figures 1D and S5). pEasyG3-derived PCR products lack sequences required for *E. coli* propagation and therefore are shorter than those generated from pEasyG1 and pEasyG2 templates; the pEasyG3 series represents a streamlined platform to optimize in vivo cloning of gRNAs in *S. cerevisiae*. Interestingly, pEasyG3-derived PCR fragments (and pEasyG2 PCR products) can be recombined in different ways to form 2–6 gRNAs (Figure 1E and 1F). Alternatively, two identical gRNAs can be cloned to specify a single Cas9 target in the genome by generating PCRs from the pEasyG3-mic and pEasyG3-zeo/nat/hph with the same gA1/gB1 primer pair (Figure 2C).

**Figure 2.**
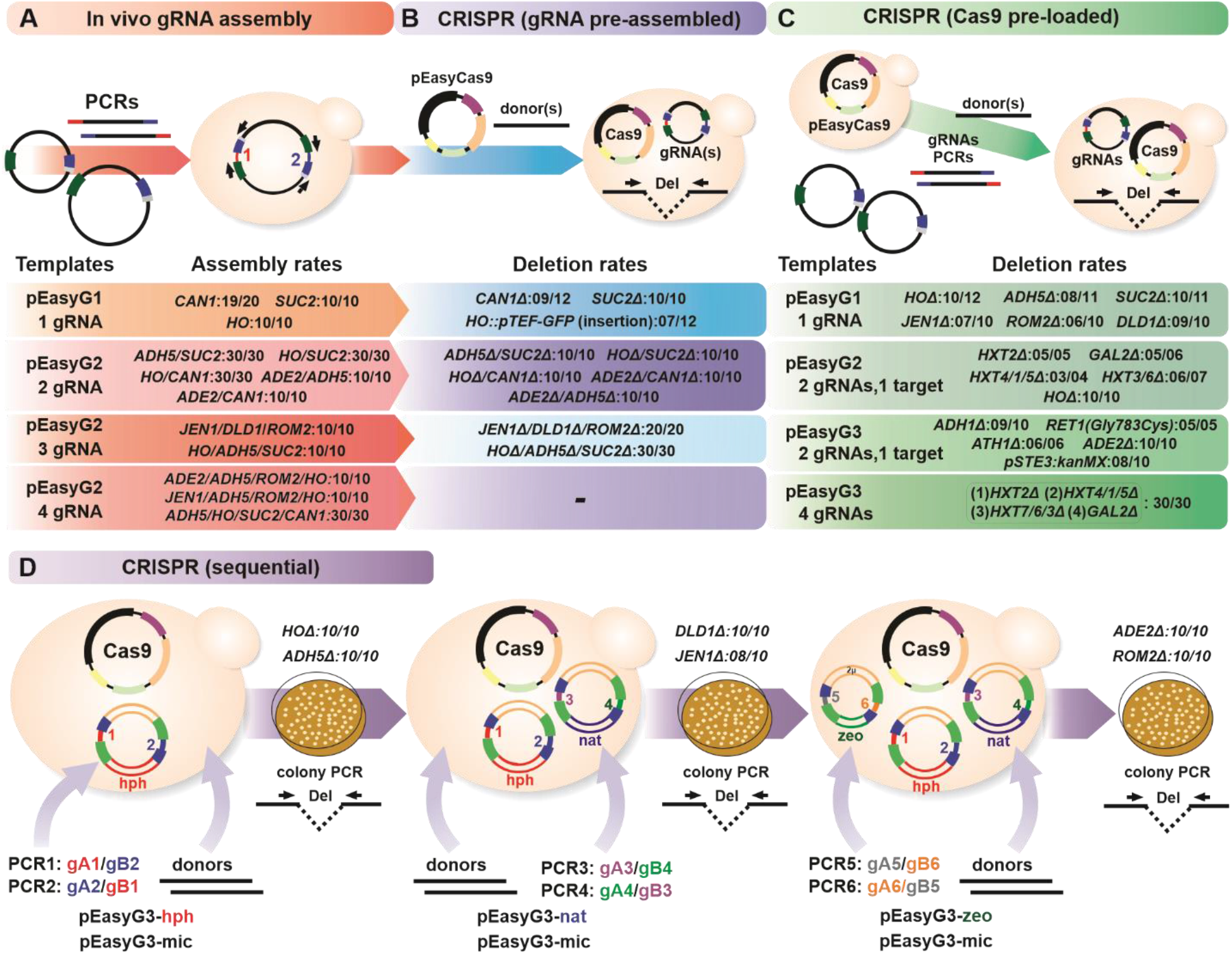
Testing the EasyGuide efficiency. (A) In vivo assembly rates (positive colonies/tested colonies) for pEasyG1/G2-derived amplicons. Correct assembly is diagnosed by colony PCRs with primers flanking the junctions between parts. The scrutiny of 30 colonies indicates three independent transformation replicates of 10 tested colonies. (B) Deletion rates were obtained by transforming strains carrying pre-assembled gRNAs with pEasyCas9 and specific donor(s). Deletion efficiency is assessed by PCR with specific flanking primers. (C) Deletion rates are tested under the Cas9 pre-loading mode. (D) A sequential genome editing procedure by successively transforming two gRNAs and respective donors, while maintaining the pEasyCas9. Rates obtained for the sextuple deletion *HOΔ, ADH5Δ, JENΔ, DLD1Δ, ADE2Δ, ROM2Δ* are shown. Schemes for PCR primer combinations and agarose gel electrophoreses for all experiments are shown in Figures S2-S20.

**Testing the EasyGuide efficiency**

We first tested the EasyGuide system during specifically designed experiments and then in our lab projects for editing genes of interest in *S. cerevisiae* (Figures 2 and S7–S20). Initially, we assessed the efficiency of *S. cerevisiae*-mediated assembly of pEasyG1 and G2-derived PCR fragments (Figure 2A). Overall, gRNA plasmid assembly in *S. cerevisiae* was highly efficient, rendering hundreds of colonies from a transformation with 10–20 µL of PCR products derived from pEasyG1 or pEasyG2(3)-mic and pEasyG2(3)-zeo/nat/hph templates (Figure S16). Correct assembly rates of 100% were observed in most cases (Figure 2A). We tested the in vivo concatenation of four pEasyG2 modules into a single replicon (i.e., four gRNAs) by analyzing five independent transformation events (three different gRNA configurations), and for each one scrutinizing 10 colonies with six PCR combinations, including the use of spacer-specific primers (see Figures S15–S18). We scored 100% positives across all 50 colonies tested, indicating that in vivo assembly of pEasyG parts is optimal when each module is selected according to its corresponding antibiotic in the medium (Figure S16).

We then tested the CRISPR/Cas9 deletion of targeted genes by transforming the pEasyCas9 and donors into strains in which the gRNAs have been pre-assembled (Figure 2B). We usually obtained high deletion rates for up to three targets for highly efficient spacers whose sequences were retrieved from the literature or designed in our lab (see materials and methods). However, the number of transformed colonies decreased dramatically for more than two gRNAs, and we repeatedly failed to obtain any colonies when deletions were attempted with four different gRNAs. We solved this problem by transforming four gRNAs and donor fragments to delete genes encoding hexose transporters into a strain carrying the pEasyCas9 plasmid (i.e., the Cas9 pre-loading mode) (Figures 2C and S19). Four to six genes can also be deleted efficiently through sequential steps (Figure 2D). First, two donors are co-transformed with pEasyG3-mic and pEasyG3-hph PCR fragments into a Cas9-expressing strain. From a propagated positive colony, a second round of transformation is performed with pEasyG3-mic and pEasyG3-nat amplicons and two further donors. In a third round, co-transformation of the fifth and sixth gRNAs and donors complete the genome editing experiment (Figures 2D and S20). It should be noted that deletions introduced by sequentially assembling gRNA-encoding plasmids were accomplished without curation of the previously transformed plasmid.

### One-day genome editing procedure

The Cas9 pre-loading mode offers a streamlined and highly efficient CRISPR/Cas9 genome editing workflow (Figure 2C and 2D). This approach yields more transformants than using pre-assembled gRNAs, regardless of the gRNA(s) and genomic target(s) selected. The gRNA assembly, gRNA expression, Cas9-mediated DNA cleavage, and donor double-strand break repair, are all coupled in the Cas9 pre-loading mode, which significantly reduces the duration of CRISPR/Cas9 genome editing procedure to a single day. First, PCR products of gRNA(s) modules are generated from pEasyG1-3 plasmids. In parallel, genomic DNA with primers specifying a genetic modification can amplify one or more donors using PCR (see materials and methods). Agarose gel electrophoresis can confirm accurate amplification of donors and gRNAs. From the reaction tubes without any PCR product cleanup, approximately 10–20 µL of pEasyG-derived fragments and 20–40 µL of donor can be co-transformed into pEasyCas9 pre-loaded cells thawed from a −80°C glycerol stock. The resulting colonies can be screened by PCR after a three-day incubation on solid medium.

## PERSPECTIVES

Here we have showed how EasyGuide plasmids can quickly and efficiently delete up to four targets at once and six targets in sequence, providing a valuable tool for routine genome editing in yeast genetics labs. By cloning mutant and chimeric forms of Cas9 fused to regulatory modules for generating modified pEasyCas9 plasmids, the portfolio of EasyGuide applications can be expanded to include CRISPR interference, activation, DNA deamination, and further CRISPR approaches.^1,16^ More than six gRNAs can be assembled into the cell by subcloning pEasyG2 and G3 modules to include additional selection marks (e.g., auxotrophic). The pEasyG3 modules are also well suited for concatenating parts through Golden Gate assembly, supporting even more complex multi-gRNA constructions.^10^ These may combine pEasyG3 parts with gRNA expression enhancers such as gRNA-tRNA array modules.^8^ The simple and versatile EasyGuide plasmid system complements the CRISPR/Cas9 toolkit, especially for yeast molecular geneticists searching for streamlined CRISPR/Cas9 applications.

## EASYGUIDE PLASMIDS AVAILABILITY

The plasmids are available at Addgene under the identification numbers: pEasyCas9 (184909), pEasyG1-zeo (184910), pEasyG1-nat (184911), pEasyG1-hph (184912), pEasyG2-zeo (184913), pEasyG2-nat (184914), pEasyG2-hph (184915), pEasyG2-mic (184916), pEasyG3-zeo (184917), pEasyG3-nat (184918), pEasyG3-hph (184919), and pEasyG3-mic (184920).

## ASSOCIATED CONTENT

### Supporting Information

The Supporting Information is available free of charge

Materials and Methods describing pEasyGuide plasmid construction; strains and growth conditions; gRNA and donor designs; and pEasyGuide-mediated CRISPR/Cas9 genome editing (PDF). Supplementary Figures S1−S20 displaying the pEasyGuide plasmids maps and agarose gel electrophoreses of the CRISPR/Cas9 experiments (PDF); Supplementary Tables S1-S3 showing oligonucleotides used in this work and primers for donor PCR amplifications (Excel).

## Author Contributions

A.P.J., L.V.S., and J.G. conceived the study and designed the experiments. A.P.J., L.S.B., and J.A.B. constructed the pEasyG plasmids. A.P.J., J.A.B., Y.A.M., L.S.B., I.F., and J.G.R.B. performed the experiments. I.F., J.G.R.B., and L.V.S. established the Cas9 pre-loading approach; A.P.J., L.V.S., and J.G. wrote the manuscript. All authors critically revised the manuscript.

## ACKNOWLEDGMENTS

This work has been supported by the São Paulo Research Foundation grants FAPESP 2017/13972-1 to J.G., FAPESP 2017/24453-5 to A.P.J., and FAPESP 2020/07918-7 and Serrapilheira Institute (Serra-1708-16205) to L.V.S.

