## Supplementary material for "EasyGuide plasmids support in vivo assembly of gRNAs for CRISPR/Cas9 applications in *Saccharomyces cerevisiae*": Materials and Methods

**pEasyGuide plasmids construction.** The pEasyCas9 plasmid, constitutively expressing the *S. pyogenes* Cas9 codon-optimized for *S. cerevisiae*, was subcloned from the pDuRCC plasmid.[1] Two PCR fragments of 3,152 bp and 5,248 bp were amplified from pDuRCC with primer pairs Ori_f3 / 2micron_r, and MXterm_f2 / CYCterm-r, respectively (Table S1). The kanMX cassette was amplified from the pUG6 plasmid[2] with oligos MX5_f / MXterm_5r (Table S1). The three fragments were assembled in vivo in *E. coli*[3] generating a 9,549 bp plasmid (Figure S1).

The pEasyG1 plasmids were constructed in a stepwise manner. First, a PCR fragment of 2,737 bp, harboring the 2micron origin, pSNR52, and the gRNA scaffold, was generated from the pDuRCC plasmid with primers Amp_f / PromMX2_r. In parallel, a 2,511 bp fragment containing the zeoMX cassette was amplified from the pUG66[4] with primers PromMX2-f / Amp_r. The two overlapping fragments were fused in vivo in *E. coli*[3] generating the plasmid pJASPR3-zeo (data not shown). The pJASPR3 was linearized by a PCR (5,045 bp) with primers JACa_Tn5 / JACb_Tn5 and recombined into *E. coli* with a 1,578 bp fragment amplified from the pKD13[5] with oligonucleotides pKD13f-1 / pKD13r-1. The pKD13-derived fragment was inserted in between the promoter pSNR52 and the gRNA scaffold, providing a cloning placeholder and a kanamycin resistance marker. The resulting pEasyG1-zeo plasmid has 6,538 bp (Figure S2A). The pEasyG1-nat (6,730 bp) and pEasyG1-hph (7,186 bp) were generated in yeast through in vivo exchange of the MX cassette via homologous recombination with amplicons generated with primers PromMX10f / MXterm_5r from pAG25 (natMX) and pAG34 (hphMX), respectively.[6] After antibiotic selection with nourseothricin or hygromycin, the resulting plasmids were purified from *S. cerevisiae* and transformed into *E. coli* by electroporation.[7]

PCR fragments of 3,983 bp, 4,175 bp, and 4,631 bp were respectively generated from pEasyG1-zeo/nat/hph with primers O_f / Scaf_r. A 32-nts overlapping between the amplicon extremities ensured the *E. coli* in vivo circularization of each PCR fragment to respectively generate the plasmids pEasyG2-zeo/nat/hph (Figure S3A). The pEasyG2-mic was cloned in two steps. First, the pUC19 vector[8] was linearized by PCR with primers Vf / Vr (1,725 bp). Two additional fragments were generated from the pEasyG1 with primer pairs 2gScaf_f / 2gTer_r (137 bp), and 2gProm_f / 2gProm_r (314 bp). The three fragments were assembled via circular polymerase extension cloning (CPEC)[9] resulting in the plasmid p2gRNA (2,093 bp, data not shown). This plasmid was a template to generate a 946 bp PCR fragment with primers SNRamp / OriJACa. This fragment was combined, through *E. coli* in vivo cloning, with a 2,748 bp PCR fragment, amplified from pEasyG1, to generate the 3,635 bp pEasyG2-mic (Figure S3B).

Plasmids of the pEasyG3 series were assembled through the Golden Gate method.[10] PCR fragments of 1,350, 1,548, and 2,005 bp were generated from the pEasyG2-zeo/nat/hph, respectively, with the primers MX_fwd / SNR52_rev. Another amplicon of 228 bp encoding the gRNA scaffold was produced from the pEasyG2 with primers gRNA_frd / gRNA_rev. The two PCR fragment types were ligated in vitro with the pGGA vector according to the manufacturer's instructions (NEBridge® Golden Gate Assembly Kit, BsaI-HF® v2; New England BioLabs, Inc) and transformed into *E. coli*. The resulting pEasyG3-zeo/nat/hph plasmids (Figure S5A) encode a chloramphenicol resistance marker for selection in *E. coli*. The pEasyG3-mic (Figure S5B) was also assembled from two PCR fragments ligated into the pGGA backbone (New England BioLabs, Inc). One PCR product of 1,702 was generated from the pEasyG1 with primers gRNA_fwd / 2MGold_rev. This was combined via Golden Gate assembly with a 291 bp PCR fragment amplified from the pEasyG2-mic with primers SNR52Gold_f / SNR52_rev.

Oligonucleotides used for Golden Gate assemblies were designed with the assistance of the NEBridge Golden Gate Assembly Tool 2.6.0 (https://goldengate.neb.com/#!/). Primers used for *E. coli* in vivo cloning and CPEC were manually designed to contain a minimum of 20-nts overlapping between the connecting parts. Oligonucleotides were synthesized at Exxtend Biotecnologia Ltda (Campinas, São Paulo, Brazil). The complete list of oligonucleotides used in this study is available in Tables S1 and S2. PCRs were performed with Phusion® High-Fidelity DNA Polymerase using standard HF buffer following the manufacturer's protocol (New England BioLabs, Inc). An annealing temperature of 58 °C and 72 °C extension for 15 sec per 500 bp were set for most PCRs. For templates that are difficult to amplify different primer annealing temperatures were tested and PCRs were performed with Phusion GC buffer. The resulting PCRs fragments were purified with the QIAquick PCR Purification Kit or with the QIAquick Gel Extraction Kit (QIAGEN). For *E. coli* in vivo cloning, equimolar amounts of PCR fragments were mixed as described[3] and transformed into the *E. coli* strain DH5α[11] by a 42 °C thermal shock procedure.[7] Chemically competent cells were prepared with the rubidium chloride protocol.[7] Colonies resulting from the transformation were screened by PCR[3] using the GoTaq® DNA polymerase (Promega) according to the manufacturer's protocol. Upon confirmation, cells were cultured overnight in LB medium and the plasmid was extracted by the QIAGEN Plasmid Mini Kit (QIAGEN). Key structural parts of plasmids generated in this study were confirmed by Sanger sequencing and repeatedly authenticated through the generation of fragments for in vivo cloning in *S. cerevisiae* and after their in vivo assembly into more complex plasmids.

**Strains and growth conditions.** *E. coli* DH5α cells harboring the pEasyGuide plasmids were propagated or selected at 37 °C in liquid or solid LB medium with 100 µg/mL ampicillin (pEasyG1 series, pEasyG2-mic, and pEasyCas9), 50 µg/mL kanamycin (pEasyG1 series and pEasyG2-zeo/nat/hph), or 50 µg/mL chloramphenicol (pEasyG3 series). *E. coli* cultures replicating pUC19-derived plasmids with cloned donors were propagated in LB with 100 µg/mL ampicillin. Frozen cell stocks of *E. coli* were prepared by mixing them with glycerol to a final concentration of 25% and storing 1 mL aliquots at -80 °C. The haploid *S. cerevisiae* strain S288C (*MATα SUC2 gal2 mal2 mel flo1 flo8-1 hap1 ho bio1 bio6*)[12] was used throughout this work for in vivo cloning procedures and CRISPR/Cas9 genome edits. The haploid LVY_X5, a *S. cerevisiae* PE-2 derivative strain[13], was used specifically for the sequential deletion of hexose transporters (see Figure S11). Yeast cultures were propagated at 28 °C in liquid or solid YP (1% yeast extract, 2% peptone) with 2% sucrose, or in YP 2% glucose for *SUC2Δ* strains, or YP 2% maltose for selection of transformants having *HXT2*, *GAL2*, *HXT7/6/3*, and *HXT4/1/5* deleted. For yeast liquid cultures an Innova shaker incubator (New Brunswick Scientific) was used at 28 °C, 120-200 rpm. Antibiotics were used alone or in combinations according to the pEasyGuide selection requirements. The antibiotics concentrations of use are: 200 µg/mL geneticin, 250 µg/mL zeocin, 100 µg/mL nourseothricin, and 300 µg/mL hygromycin. For cryopreservation, yeast cells collected after overnight growth were mixed with glycerol to a final concentration of 25% v/v and stored at -80 °C.

**gRNA and donor designs.** A complete list of gRNA spacers and corresponding gA and gB oligonucleotides used in this study is available in Table S2. The *CAN1*[14], *ADE2*[14], and *ADH5*[15] gRNA spacers were retrieved from the literature. Other gRNAs were designed with the assistance of CRISPOR software version 4.99[16] or manually scanned upstream of Cas9 PAM sequences on the targeted loci. In the latter case, the selected 20-nts sequence was matched against a comprehensive list of spacer sequences predicted to have no off-target effects on the S288C genome.[17] The spacer sequences were specified at the 5’ end of oligonucleotides in fusion with the core gA or gB sequences (see Figure 1B, Table S2). Longer versions of gA and gB core sequences were used to design most pEasyG oligonucleotides for this work (see Table S2). However, a shorter version of gA (‘5-N(20)GTTTTAGAGCTAGAAATAGCAAGTT-3’) and gB (‘5-N(20)GATCATTTATCTTTCACTGCGGAGAAG-3’) also enables optimal amplification of pEasyG PCR fragments at an annealing temperature of 60 °C.

DNA donor sequences and templates to repair Cas9-induced double-strand breaks and introduce genome edits were generated by PCR through different approaches. Donor specifications for each target locus are listed in Table S3. First, DNA deletions were specified within bipartite oligonucleotides. These were designed to contain a 5’ region with 40-nts homology annealing upstream of the planned deletion sequence, while a 3’ region of 23-26-nts defined a core primer annealing downstream of the deleted region (i.e., on the other side of the deletion). A PCR was performed with this oligonucleotide in combination with a reverse primer biding distally downstream to the target. Once a deletion was already established on the yeast genome, a donor could be simply generated by a PCR with primers flanking the deletion site. For this purpose, genomic DNA of a deleted strain was prepared through cell mechanical disruption (zirconium beads), phenol/chloroform purification, RNAse treatment (4 µL of 100 mg/mL for 15 min at 37 °C), and ethanol precipitation. Donors were also generated for inserting two overlapping PCR fragments into the *HO* locus to assemble a gene-expressing construct. By this mean, a PCR fragment of 409 bp, containing the *TEF2* promoter from *Ashbya gossypii* (TefGFP_f / TefGFP_r)[2], was fused to the 1,050 bp GFP encoding cassette[18], which was PCR amplified with GFP_f / TDH1_r primers (Figure S7C). Similarly, the construct *pSTE3-hphMX* was assembled into the *HO* locus (Figure S14E) from PCR fragments *pSTE3* (940 bp) and *kanMX* (1,286 bp) (Table S3). In a different approach, the 316 bp donor *RET1* was designed to specify SNPs leading to the amino acid Gly783Cys exchange at the encoded polypeptide. In addition, the introduced SNPs enabled the design of a derived CAPs marker[19] through PCR amplification of the edited DNA with the primers RET1SalI_f / RET1SalI_r and cleavage of the product with the SalI restriction enzyme (Figure S14C).

In some cases, donor sequences specifying targeted deletions were pre-cloned in *E. coli*. The pSUC2donor plasmid was generated from two genomic PCR products of 315 bp (SucDelL_f / SucDelL_r) and 442 bp (SucDelR_f / SucDelR_r) assembled via CPEC[9] into a PCR linearized pUC19 plasmid (primers Vf / Vr, 1,725 bp). Similarly, pADHO was assembled via CPEC from fragments of 410 bp (DonAHO_f / AHO_rev), and 387pb (AHO_for / DonAHO_r) amplified from the genome of *ADH5Δ* and *HOΔ* deleted strains, respectively, and fused to the 1,725 bp pUC19 amplified backbone (Vf / Vr). A fused *ADH5Δ-HOΔ* donor of 753 bp generated by PCR from the pADHO (DonAHO_f / DonAHO_r) was used to simultaneously delete the two targeted genes on the genome (Figure 2D). The donors used for one-step deletion of hexose transporters (Figure S19) were pre-assembled in vivo into *S. cerevisiae* S288C. The pAB_DonGAL2/HXT763 was assembled from seven PCR fragments: (1) gA_HXT3 / gB_GAL2 (3,101 bp from pEasyG3-hph); (2) gB_HXT3 / VrHXT3 (1,446 pb, pEasyG2-mic); (3) gA_GAL2 / 2MicGAL2 (1,611 pb, pEasyG2-mic); (4) Gal2P_f / Gal2P_r (459 pb, S288C genomic DNA); Gal2T_f / Gal2T_r (483 pb, genomic DNA); (6) Hxt7P_f / Hxt7P_r (444 pb, genomic DNA); (7) Hxt3T_f / Hxt3T_r (454 pb, genomic DNA). A fused *GAL2Δ-HXT763Δ* donor of 1,014 bp was PCR amplified with Gal2P_f / HXT3ter_r from the pAB_DonGAL2/HXT763 and used to delete the two loci (Figure S19C and S19D). The pAB_DonHXT2/HXT415 was also assembled from seven PCR fragments: (1) gA_HXT2 / gB_HXT14 (2,645 bp from pEasyG2-nat); (2) gB_HXT2 / VrHXT5 (1446 pb, pEasyG2-mic); (3) gA_HXT14 / 2MicHXT2 (1611 pb, pEasyG2-mic); (4) HXT2P_f / HXT2P_r (439 pb, S288C genomic DNA); (5) HXT2T_f / HXT2T_r (389 pb, genomic DNA); (6) HXT4P_f / HXT4P_r (435 pb, genomic DNA); (7) Hxt5T_f / Hxt5T_r (517 pb, genomic DNA). A fused *HXT2Δ-HXT4/1/5Δ* donor of 964 bp was PCR amplified with primers HXT2pro_f / HXT5term_r from pAB_DonHXT2/HXT415 and used to simultaneously delete the two loci (Figure S19A and S19B).

**pEasyGuide-mediated CRISPR/Cas9 genome editing.** PCRs to amplify the pEasyG parts were performed with the Phusion® High-Fidelity DNA Polymerase with the GC buffer according to the manufacturer's instructions (New England BioLabs, Inc). pEasyG amounts ranging from 20-60 ng were used as PCR templates in a 50 µL reaction mix containing a specific gA/gB primer pair (Table S2). PCR settings were: 98 °C denaturation (15 sec), an annealing temperature of 60

°C (20 sec), and 72 °C extension for 15 sec per 500 bp; 35 PCR cycles. The resulting pEasyG amplicon sizes are as described in the main text. For CRISPR/Cas9 experiments, a donor(s) specifying the genome edit was generated in parallel via PCR under similar conditions using the Phusion HF buffer. After PCR, electrophoresis on an 2% agarose gel stained with ethidium bromide was performed to confirm the pEasyG and donor amplifications. Transformations of pEasyG amplicons and donors into S288C cells were conducted using the LiAc/SS carrier DNA/PEG protocol as described.[20]

Throughout this work, two approaches were taken for CRISPR/Cas9 genome edition (Figure 1A). Under a Cas9 pre-loaded mode, we first transformed the pEasyCas9 plasmid into the S288C strain. After confirmation of transformation and propagation of transformants (28 °C, 210 rpm), cells were collected at an $OD_{600}$ ~0.6, pelleted, washed, and resuspended for aliquoting 100 µL of cells with 5% v/v glycerol and 10% DMSO. Cells were stored at -80 °C.[21] Yeast cell aliquots were thawed for use and transformed with about 10-20 µL of the resulting pEasyG PCR products and 20-40 µL donors taken directly from the respective PCR tubes, or with about 1 µg of a silica column purified donor. Only in the case of pEasyG1-derived amplicons, treatment with 10 U DpnI (New England BioLabs, Inc.) for one hour is necessary to preclude template carry over leading to false-positive transformants. For transformation of more than two gRNAs (i.e., when more than two genome loci were targeted), pEasyG and donors PCR products were purified (QIAquick PCR Purification Kit, QIAGEN), vacuum concentrated (RVC 2-18 CDplus, Martin Christ Freeze Dryers), and resuspended with water for the use of 1-5 µg of each PCR product in the transformation. Following a 45 min 42 °C thermal shock, cells were recovered for one hour in 15 mL YPD 2% sucrose inside 250 mL shake flasks at 210 rpm. After outgrowth, cells were spun down, resuspended in 100 µL YPD 2%, and plated onto selective medium with geneticin (pEasyCas9) and zeocin (or nourseothricin/hygromycin) according to the pEasyG backbone(s) used. After three days of growth at 28 °C, DNA was prepared by transferring colony cells with a pipette tip into 30 µL water and heating the cell suspension at 96 °C in a hot block for 10 min. For each sampled colony DNA, 12.5 µL of the prep was added to an 8.5 µL PCR mix with 1 U Taq DNA polymerase (GoTaq®, Promega). PCR was run at 58 °C annealing temperature, 15 sec extension at 72 °C, and 35 cycles. 2% agarose gel electrophoreses images of the screenings and specific PCR primers used for checking genome edits are shown in Figures S2-S20. Oligonucleotides used are listed in Table S1.

An alternative approach was to assemble the gRNA(s) prior to the genome edit experiment (i.e., the gRNA pre-assembling mode, Figure 1A). In this case, 5-10 µL of each pEasyG PCR product was transformed into S288C cells.[20] After three days, colonies were scrutinized by PCR with specific primers to confirm the structure of assembled Yeast pEasyG replicons (pYEasyG, see Figures S2, S4, and S6). A positive colony was inoculated and propagated overnight in YPD 2% with the antibiotic necessary to maintain the pYEasyG replicon. In the next day, a 2 mL inoculum was transferred into 50 mL YPD 2% (plus antibiotic) and propagated until an $OD_{600}$ ~0.6, when cells were pelleted down and resuspended in sterile water. About 1-5 µg of each donor and pEasyCas9 PCR product are added to 100 µL cells and transformed according to the LiAc method.[20] The mixture is subjected to a 45 min 42 °C thermal shock, outgrowth for one hour at 28 °C , and then plated onto selective solid medium with geneticin for pEasyCas9 selection and the antibiotic required for pEasyG selection. After three days of 28 °C incubation, the resulting colonies were scrutinized by PCR as described above. The correspondence of the obtained PCR product for each targeted loci and the planned deleted sequence was confirmed by Sanger DNA sequencing.
