## Supplementary Figures S1-S20 for "EasyGuide plasmids support in vivo assembly of gRNAs for CRISPR/Cas9 applications in *Saccharomyces cerevisiae*"

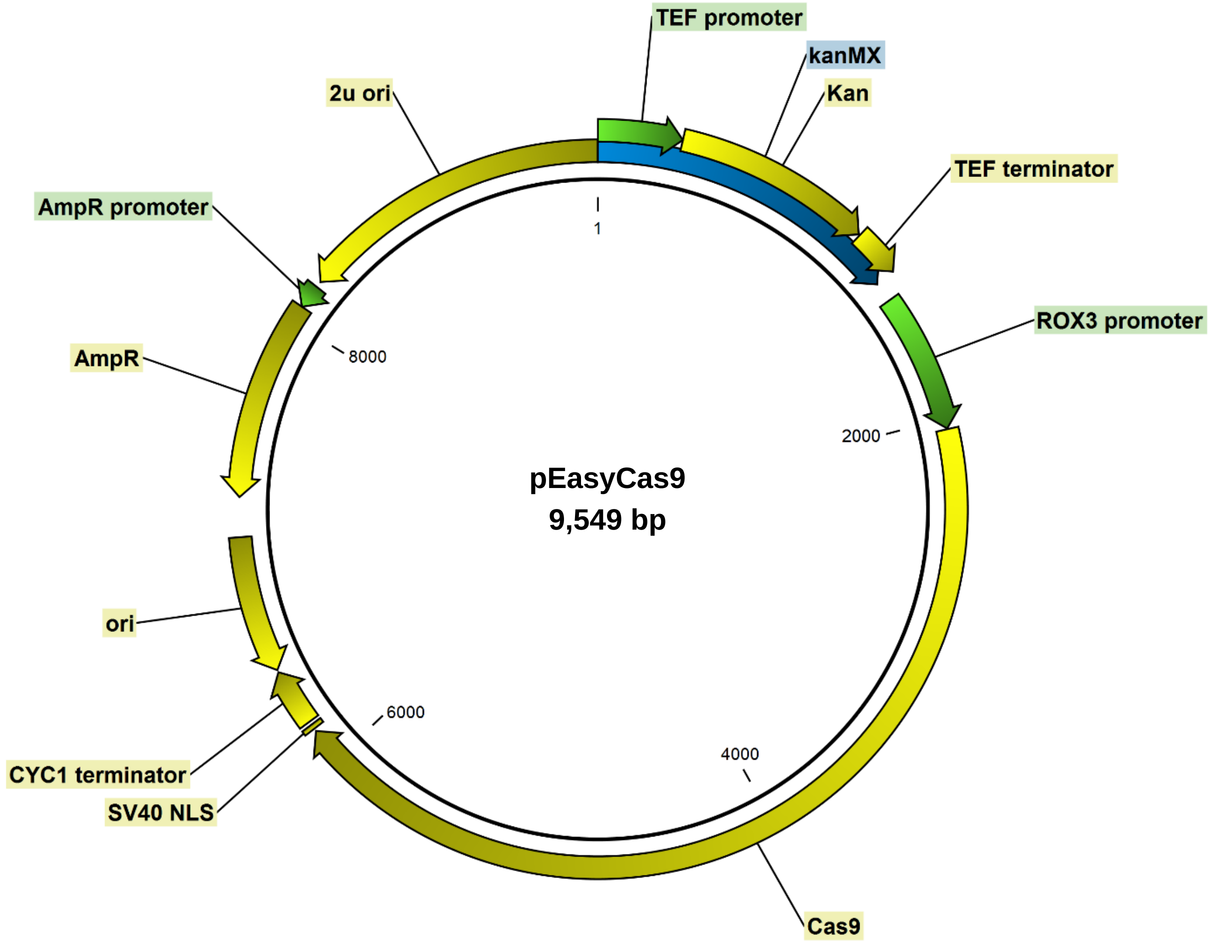


**Figure S1.** Map of pEasyCas9 plasmid.

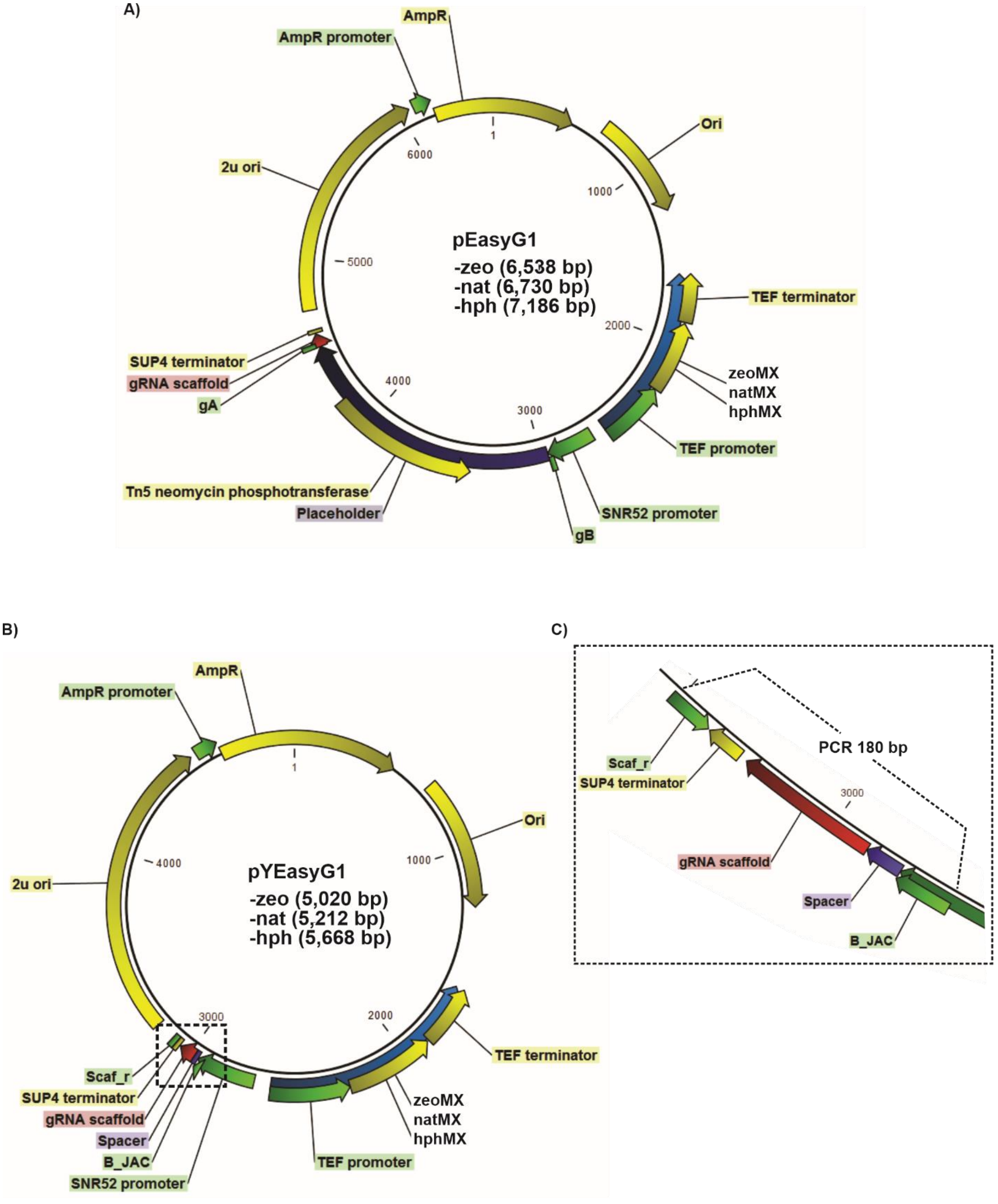


**Figure S2.** pEasyG1 plasmid. (A) A Physical map of pEasyG1 is shown. (B) The pYEasyG1 plasmid results from in vivo (*S. cerevisiae*) circularization of the pEasyG1-derived PCR fragment. The dotted square indicates the assembled gRNA structure. (C) Zoom into the gRNA structure with a 20-nts spacer resulting from the homologous recombination. A PCR of 180 bp (primers B_JAC / Scaf_r) allows for a validation of the assembly.

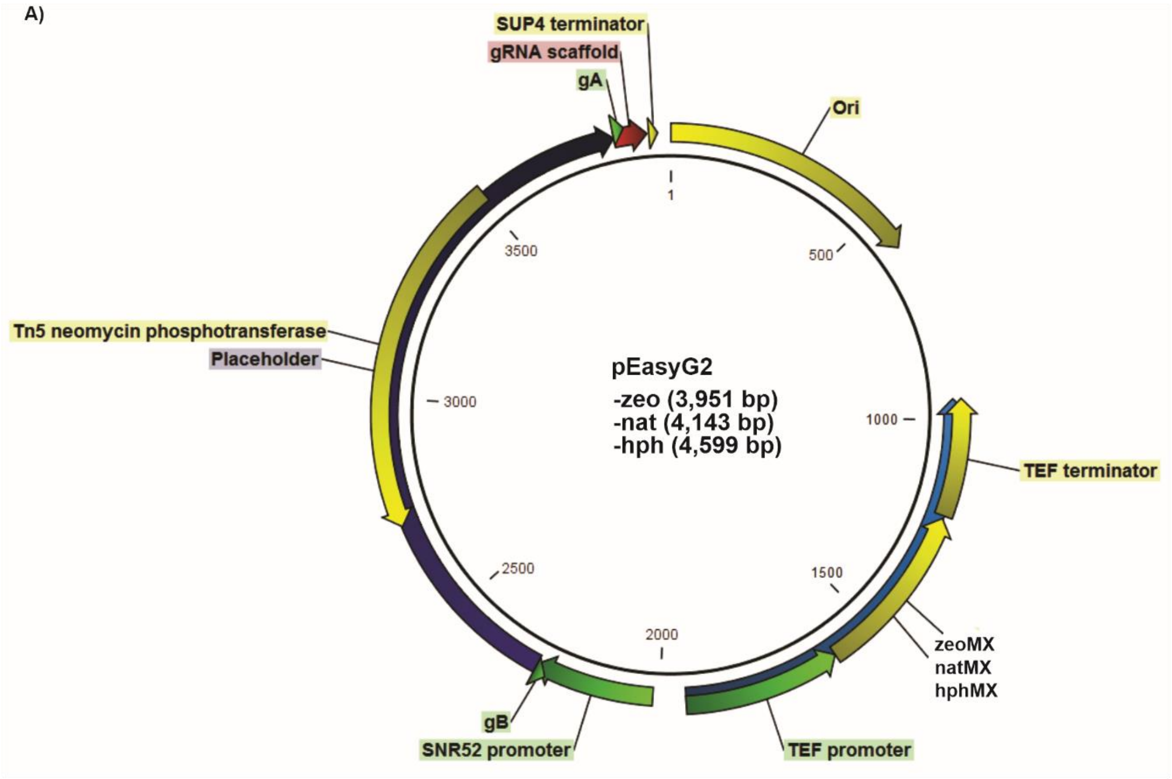


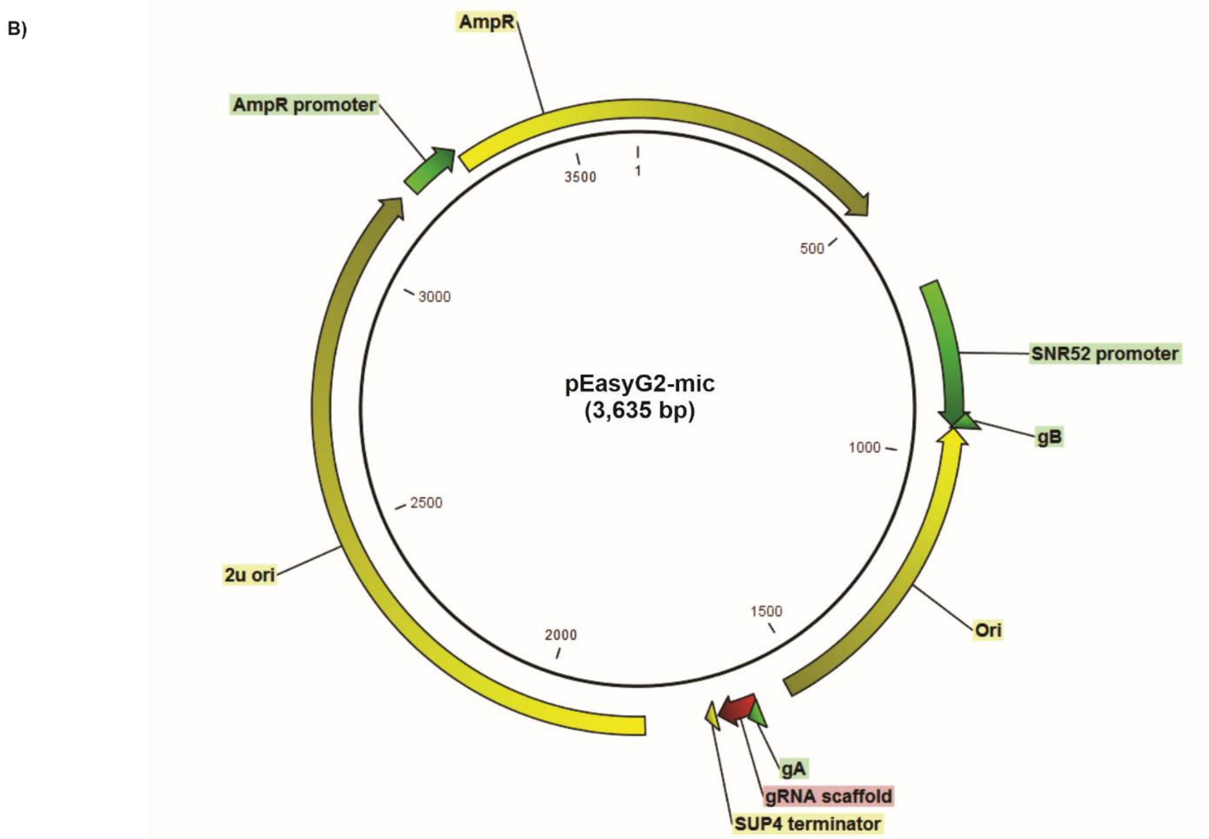


**Figure S3.** The pEasyG2 series. (A) A physical map of pEasyG2-zeo/nat/hph. (B) Map of pEasyG2-mic.

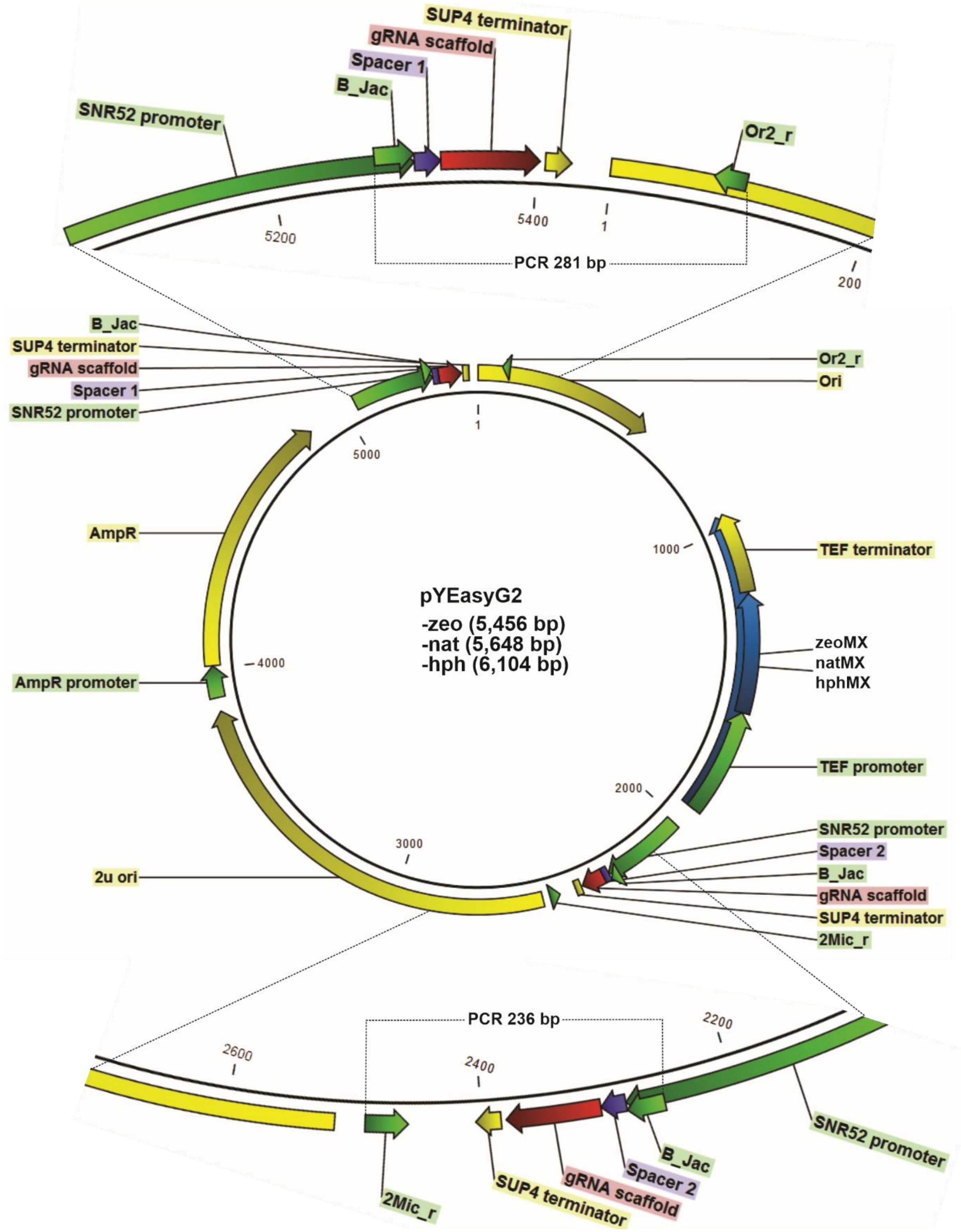


**Figure S4.** pYEasyG2 after in vivo assembly. Plasmid resulting from the recombination between pEasyG2-zeo/nat/hph and pEasyG2_mic PCR products. Structures of the two assembled gRNAs are enlarged showing PCR primers combinations to authenticate the proper connection between parts.

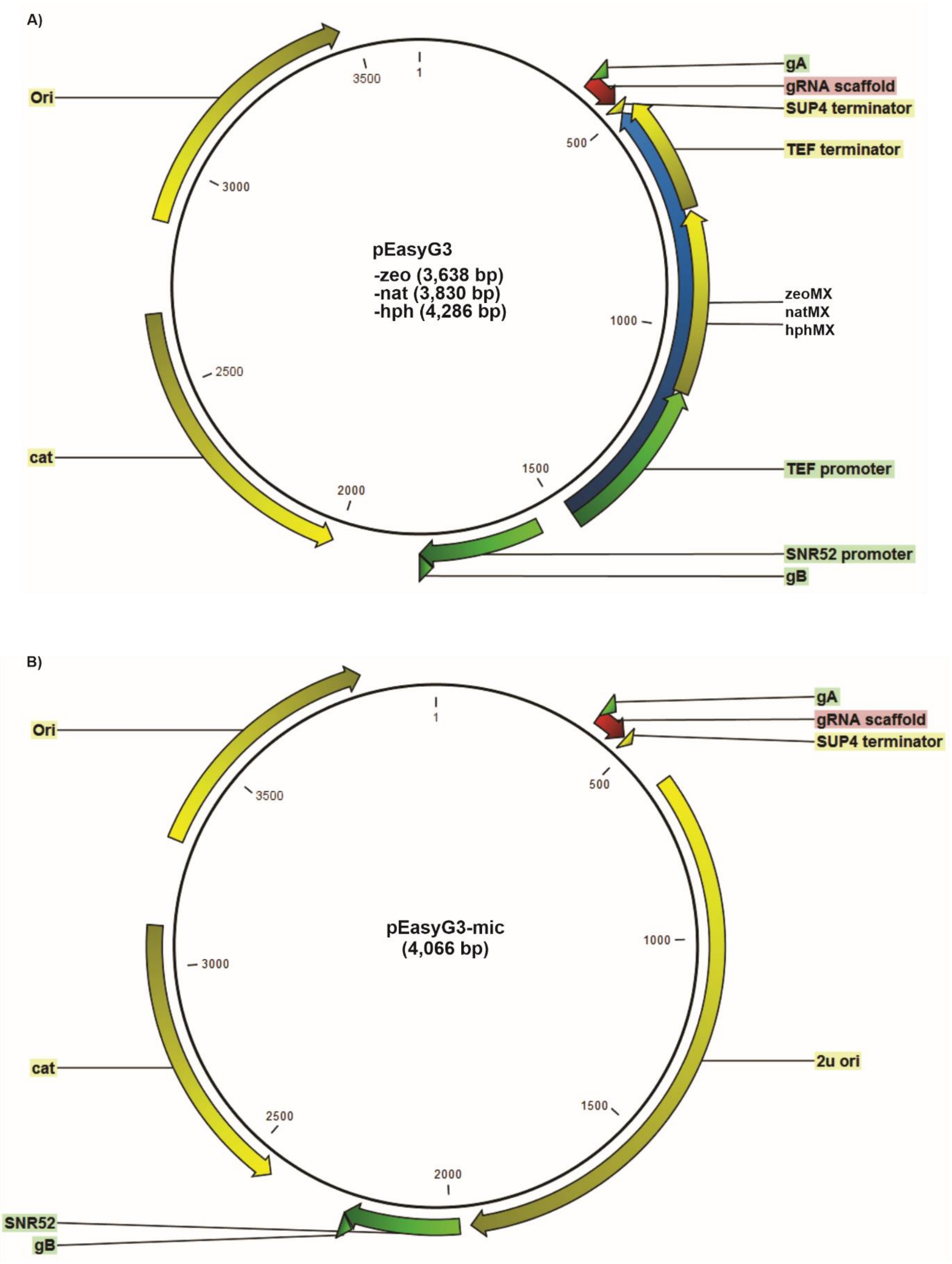


**Figure S5.** The pEasyG3 series. (A) Physical map of pEasyG3-zeo/nat/hph. (B) Map of pEasyG3-mic.

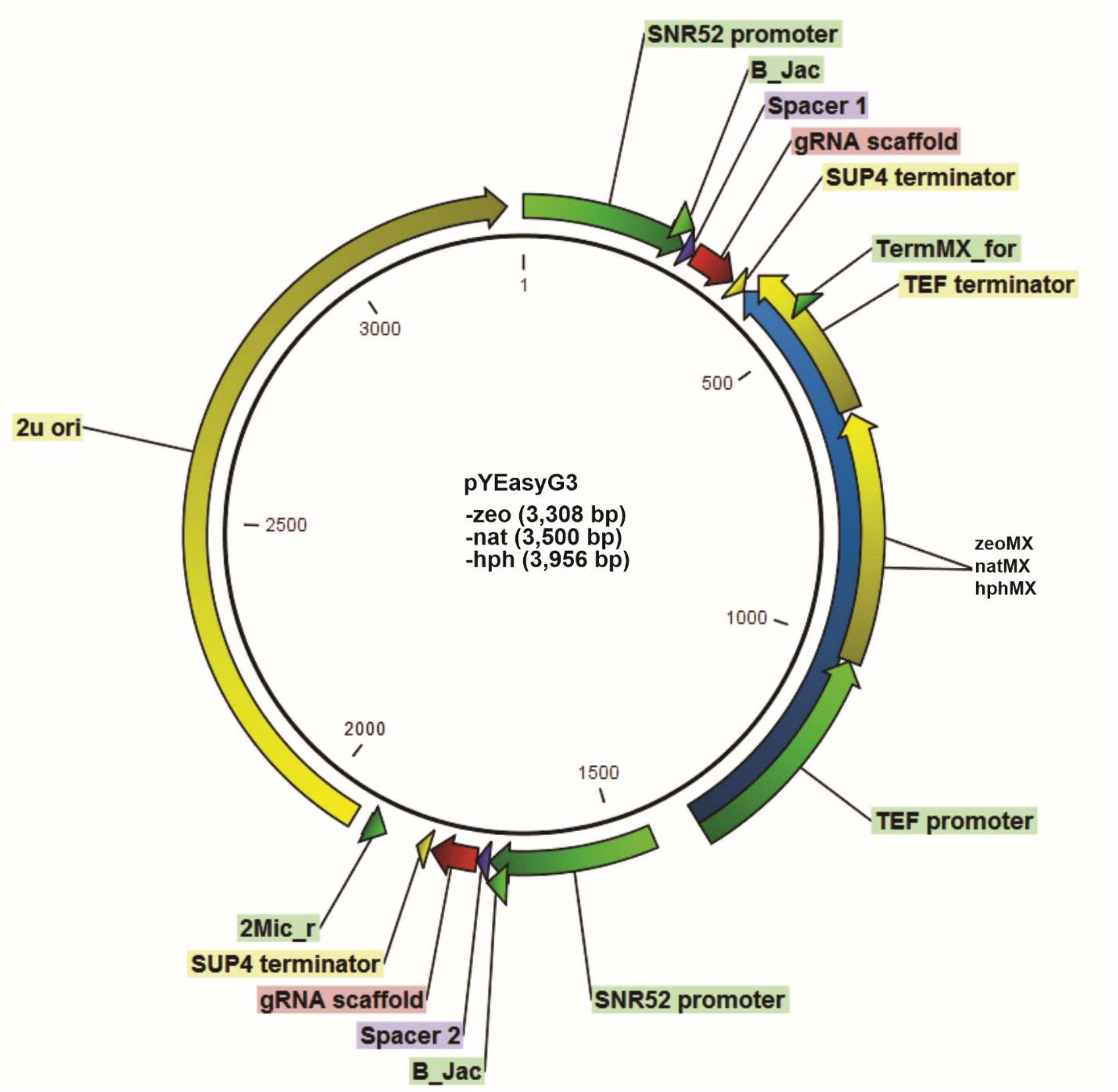


**Figure S6.** pYEasyG3 after in vivo assembly. Plasmid resulting from the recombination between pEasyG3-zeo/nat/hph and pEasyG3_mic PCR products. Two gRNAs are assembled and can be confirmed by PCR with primers combinations B_JAC / 2Mic_r (236 bp) and B_JAC / TermMX_for (232 bp).

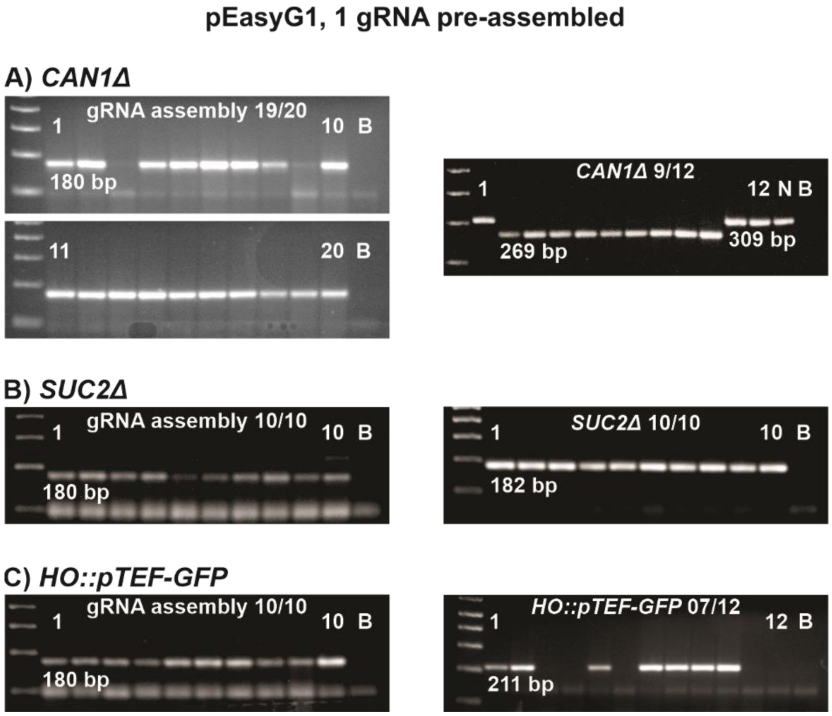


**Figure S7.** pEasyG1, 1 gRNA pre-assembled and subsequent CRISPR experiment. 2% agarose gel electrophoreses of PCR fragments are shown to estimate the rates of in vivo gRNA assembly (left) and target chromosomal modifications (right). PCR blank controls (B) were included in the agarose gels. A 100 bp ladder was used as a molecular weight standard. PCR products of 180 bp (primers B_JAC / Scaf_r, Figure S2C) are consistent with the proper in vivo circularization of pEasyG1 amplicons. The absence of a PCR product is reckoned as a negative result. 10-20 colonies were screened from each transformation. (A) *CAN1* gRNA in vivo assembly rates (left) followed by the CRISPR experiment (right) rendering nine positive deletions (269 bp PCR product with primers CAN1f2 / CAN1r2) out of 12 colonies tested. Three transformants had the same pattern as the negative control (N, a 309 bp PCR product of the wild-type locus with the same primers). (B) *SUC2* gRNA in vivo assembly rates (left) followed by the CRISPR experiment (right) showing 10 positive deletions (182 bp PCR product with primers SUC2_extL / SucTestDel_r) out of 12 colonies tested. (C) *HO* gRNA in vivo assembly rates (left) followed by the CRISPR-mediated insertion of fragments *pTEF* (409 bp) and *GFP-tTDH1* (1,050 bp) into the *HO* locus (right). A PCR product of 211 bp (primers HOEf / ProMX_rev) indicates the assembly of the two fragments and the insertion into the *HO* locus. Positive colonies had a GFP-expressing phenotype observed under the fluorescence microscope.

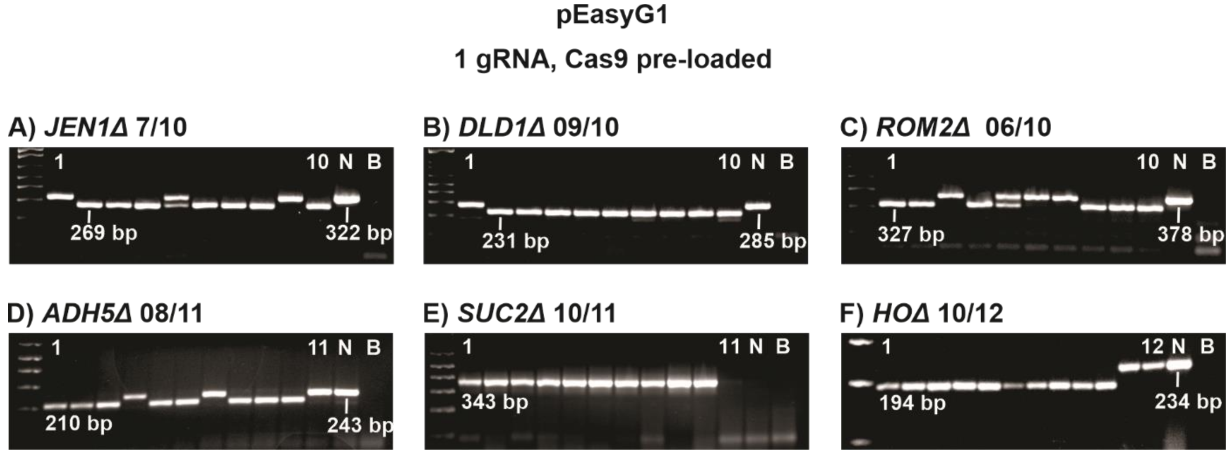


**Figure S8.** pEasyG1, 1 gRNA. CRISPR experiment with pre-loaded Cas9. 2% agarose gel electrophoreses of PCR fragments are shown to estimate the positivity of targeted deletions. Negative controls (N) of deletion, corresponding to a PCR product of the wild-type locus, and PCR blank controls (B) were loaded in the agarose gels. A 100 bp ladder was used as a molecular weight standard. 10-12 colonies were screened from each transformation. (A) *JENΔ* deletion. 269 bp PCR products from *JEN1* (primers DonJEN1_f / CheckJEN1_r) confirm seven positive deletions. A PCR product of 322 bp corresponds to an intact WT locus. (B) *DLD1Δ* deletion. 231 bp PCR products from *DLD1* (primers CheckDLD1_f1 / DonDLD1_r) confirm nine positive deletions. A PCR product of 285 bp corresponds to an intact WT locus. (C) *ROM2Δ* deletion. 327 bp PCR products from *ROM2* (primers ROM2P3for / ROM2P3rev) confirm six positive deletions. A PCR product of 378 bp corresponds to an intact WT locus. (D) *ADH5Δ* deletion. 210 bp PCR products from *ADH5* (primers ADH5f2 / ADH5r2) confirm eight positive deletions. A PCR product of 243 bp corresponds to an intact WT locus. (E) *SUC2Δ* deletion. 342 bp PCR products from *SUC2* (primers SucDelL_f / SucTestDel_r) confirm ten positive deletions. The absence of a PCR product is expected for a WT locus. (F) *HOΔ* deletion. 194 bp PCR products from *HO* (primers CheckHO_f / CheckHO_r) confirm eight positive deletions. A PCR product of 234 bp corresponds to an intact WT locus.

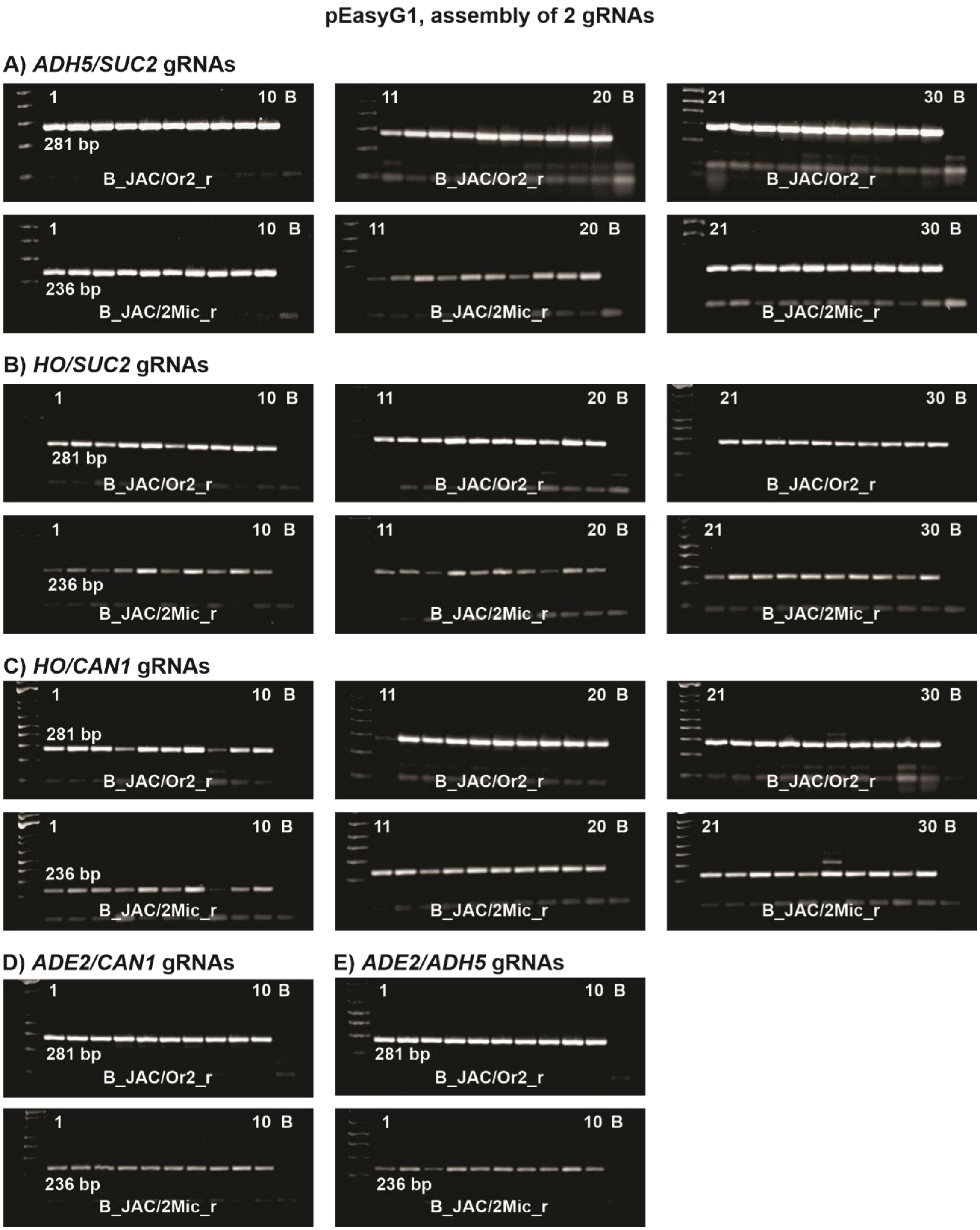


**Figure S9.** pEasyG2, in vivo assembly of 2 gRNAs. 2% agarose gel electrophoreses of PCR fragments are shown to confirm the in vivo assembly of gRNAs. PCR blank controls (B) were included in the agarose gels. A 100 bp ladder was used as a molecular weight standard. Simplified PCR screenings of colonies with the primers B_JAC / Or2_r (281 bp) and B_JAC / 2Mic_r (236 bp) show PCR products consistent with the proper assembly of the pEasyG2-hph amplicons with the pEasyG2-mic PCR product (See Figure S4). (A) Assembly of *ADH5*/*SUC2* gRNAs. (B) Assembly of *HO*/*SUC2* gRNAs. (C) Assembly of *HO*/*CAN1* gRNAs. (D) Assembly of *ADE2*/*CAN1* gRNAs. (E) Assembly of *ADE2*/*ADH5* gRNAs.

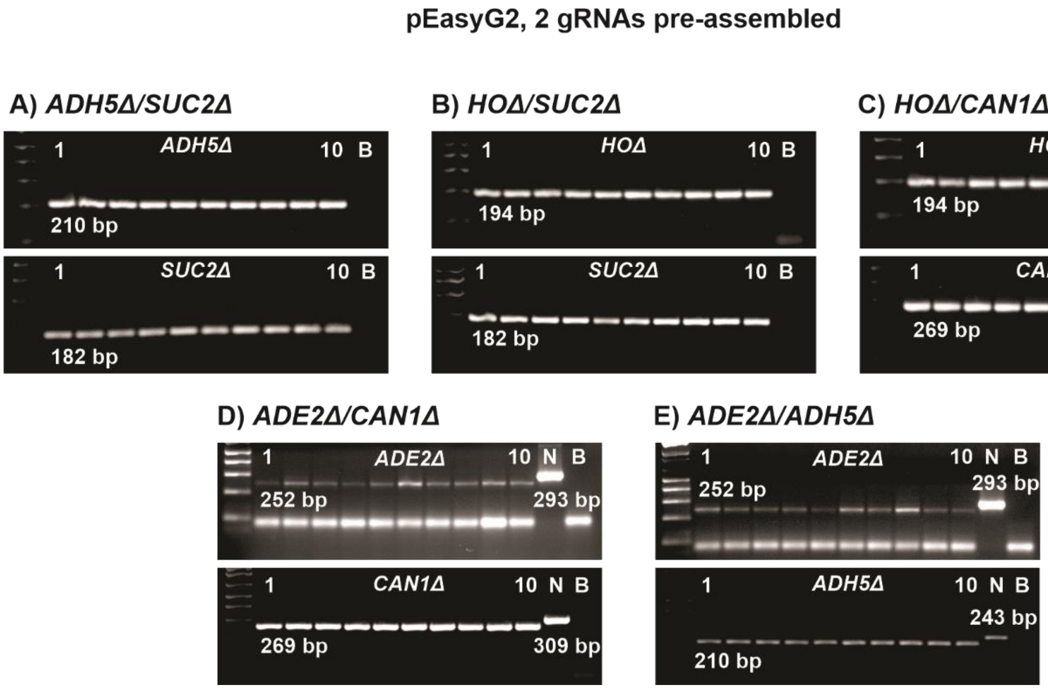


**Figure S10.** pEasyG2, 2 gRNAs for double deletions *ADH5Δ*/*SUC2Δ*, *HOΔ*/*SUC2Δ*, *HOΔ*/*CAN1Δ*, *ADE2Δ*/*CAN1Δ*, *ADE2Δ*/*ADH5Δ*. CRISPR experiment with pre-assembled gRNAs (see Figure S9). 2% agarose gel electrophoreses of PCR fragments are shown to confirm the positivity of targeted deletions. PCR blank controls (B) were included in the agarose gels. A 100 bp ladder was used as a molecular weight standard. Ten colonies from each transformation were screened. (A) *ADH5Δ*/*SUC2Δ* deletions. A 210 bp PCR product from *ADH5* (primers ADH5f2 / ADH5r2) and a 182 bp PCR product from *SUC2* (primers SUC2_extL / SucTestDel_r) confirm the deletions. (B) *HOΔ*/*SUC2Δ* deletions. A 194 bp PCR product from *HO* (primers CheckHO_f / CheckHO_r) and a 182 bp PCR product from *SUC2* (primers SUC2_extL / SucTestDel_r) confirm the deletions. (C) *HOΔ*/*CAN1Δ* deletions. A 194 bp PCR product from *HO* (primers CheckHO_f / CheckHO_r) and a 269 bp PCR product from *CAN1* (primers CAN1f2 / CAN1r2) confirm the deletions. (D) *ADE2Δ*/*CAN1Δ* deletions. A 252 bp PCR product from *ADE2* (primers ADE2f2 / ADE2r1) and a 269 bp PCR product from *CAN1* (primers CAN1f2 / CAN1r2) confirm the deletions. Negative (N) control PCRs from the WT loci are loaded (293 bp for *ADE2* and 309 bp for *CAN1*). (D) *ADE2Δ*/*ADH5Δ* deletions. A 252 bp PCR product from *ADE2* (primers ADE2f2 / ADE2r1) and a 210 bp PCR product from *ADH5* (primers ADH5f2 / ADH5r2) confirm the deletions. Negative (N) control PCRs from the WT loci are loaded (293 bp for *ADE2* and 243 bp for *ADH5*).

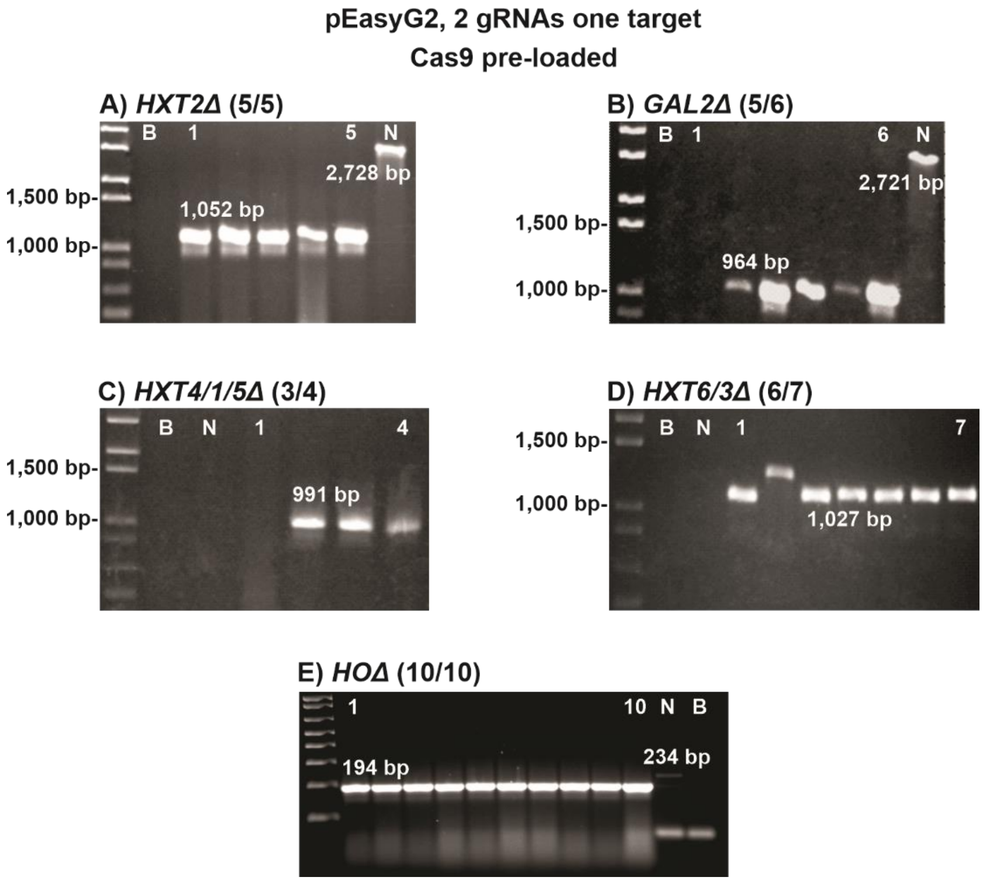


**Figure S11.** pEasyG2, CRISPR experiment using 2 gRNAs for a single deletion performed into Cas9 pre-loaded cells. 2% agarose gel electrophoreses of PCR fragments are shown from the screening of positive deletions. Four to ten colonies were tested. PCR blank controls (B) were included in the agarose gels, as well as negative (N) deletion controls consisting of a PCR with the WT DNA (not deleted) and the same primers used to screen the transformants. 1 Kb plus ladder from Thermo Scientific was the molecular weight standard. (A) *HXT2Δ* deletions. A 1,052 bp PCR product (primers FH2_f / FH2_r) confirms the deletions in 5 out of 5 colonies tested. A 2,728 bp PCR product is expected for the WT locus. (B) *GAL2Δ* deletions. A 964 bp PCR product (primers FG2_f / FG2_r) confirms deletions in 5 out of 6 colonies tested. A 2,721 bp PCR product is expected for the WT locus. (C) Deletion of the locus harboring the *HXT4, HXT1* and *HXT5* genes. A 991 bp PCR product (primers FH4_f / FH4_r) confirms deletions in 3 out of 4 colonies tested. A PCR of the WT locus (N) with the same primers would be too large to amplify (10,359 bp). (D) Deletion of the locus harboring the *HXT6* and *HXT3* genes. A 1,027 bp PCR product (primers checkDelHXT367_f / checkDelHXT367_r) confirms deletions in 6 out of 7 colonies tested. A PCR of the WT locus (N) with the same primers would be too large to amplify (6,092 bp). Experiments A-D were performed with the *S. cerevisiae* PE-2 strain and deletions were sequentially introduced in four rounds to obtain a strain with the four loci deleted. E) A 194 bp PCR product from the *HO* locus obtained with the primers CheckHO_f / CheckHO_r confirms a positive deletion in S288C. A negative (N) deletion control PCR of a WT DNA with the same primers generates a fragment of 234 bp. A 100 bp ladder was used as a molecular weight standard.

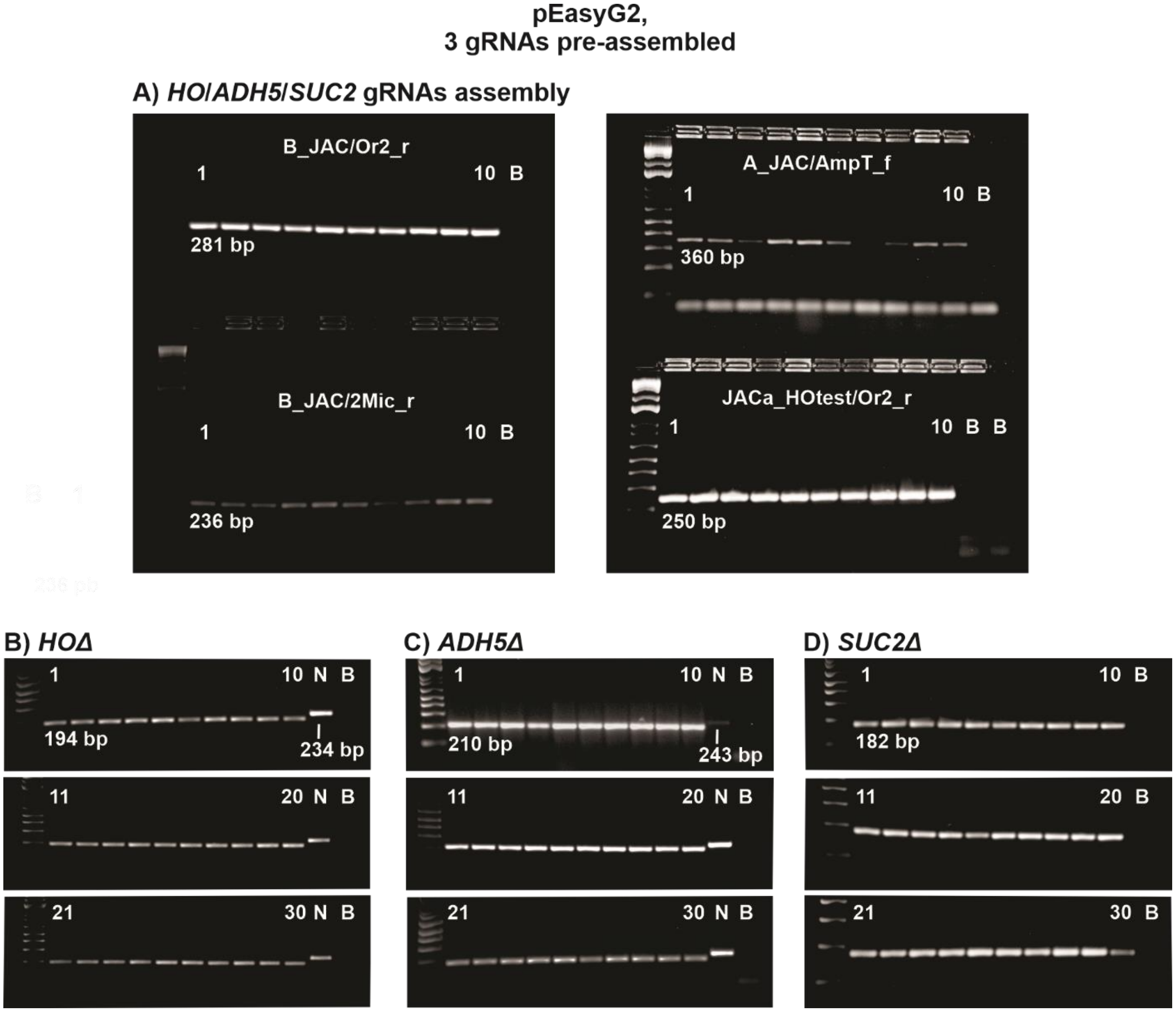


**Figure S12.** pEasyG2, 3 gRNAs for a triple deletion *HOΔ*/*ADH5Δ*/*SUC2Δ*. 2% agarose gel electrophoreses of PCR fragments are shown to confirm the in vivo assembly of gRNAs and the positivity of targeted deletions. At the PCR screening for deletions, a negative control (N), corresponding to a PCR product of the wild-type locus, and a PCR blank control (B) were included in the agarose gels. A 100 bp ladder was used as a molecular weight standard. (A) A PCR screening of 10 colonies with the primers B_JAC / Or2_r (281 bp), B_JAC / 2Mic_r (236 bp), A_JAC / AmpT_f (360 bp), and HOtest / Or2_r (250 bp) shows PCR products consistent with the proper recombination of pEasyG2-nat and pEasyG2-hph amplified modules with the pEasyG2-mic PCR product. (B) PCR screening of 30 colonies (transformation triplicates) resulting from the CRISPR experiment. A 194 bp PCR product from the *HO* locus obtained with the primers CheckHO_f / CheckHO_r confirms a positive deletion. A negative (N) control PCR of a WT DNA with the same primers generates a fragment of 234 bp. (C) A 210 bp PCR product from the *ADH5* locus obtained with the primers ADH5f2 / ADH5r2 confirms a positive deletion. A negative (N) control PCR of a WT DNA with the same primers generates a fragment of 243 bp. (D) A 182 bp PCR product from the *SUC2* locus obtained with the primers SUC2_extL / SucTestDel_r confirms a positive deletion.

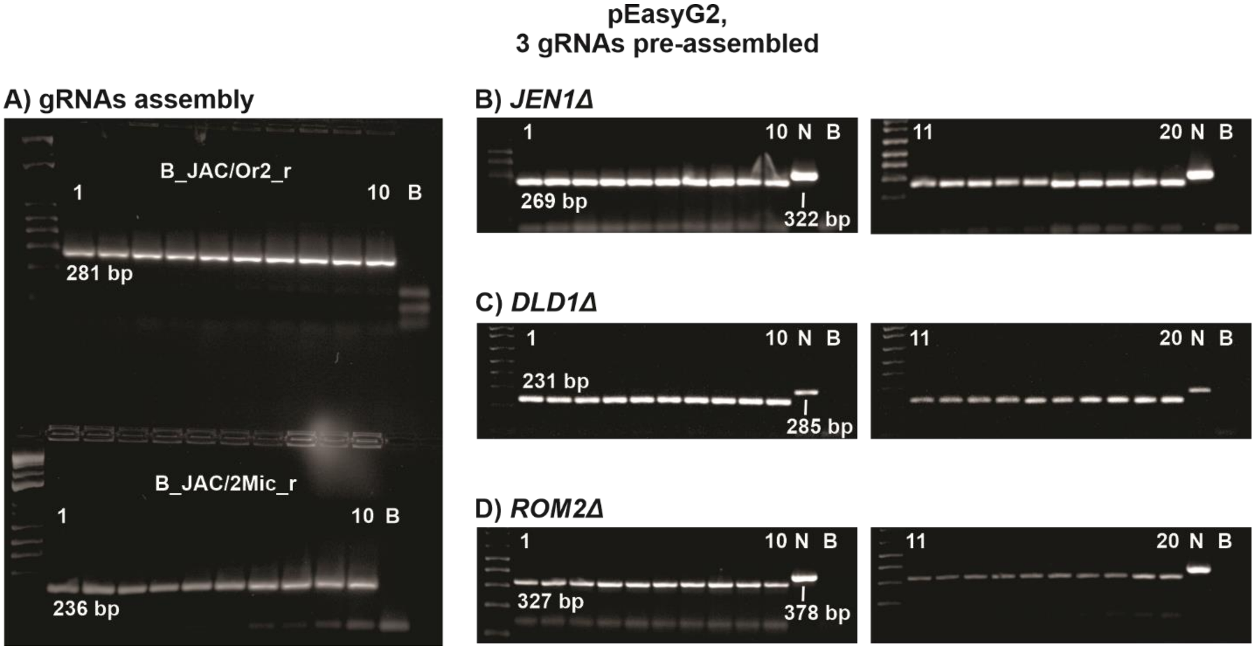


**Figure S13.** pEasyG2, 3 gRNAs for a triple deletion *JEN1Δ*/*DLD1Δ*/*ROM2Δ*. 2% agarose gel electrophoreses of PCR fragments are shown to confirm the in vivo assembly of gRNAs and the positivity of targeted deletions. At the PCR screening for deletions, a negative control (N), corresponding to a PCR product of the wild-type locus, and a PCR blank control (B) were included in the agarose gels. A 100 bp ladder was used as a molecular weight standard. (A) A simplified PCR screening of 10 colonies with the primers B_JAC / Or2_r (281 bp) and B_JAC / 2Mic_r (236 bp) shows PCR products consistent with the proper assembly of the pEasyG2-nat and pEasyG2-hph amplified modules with the pEasyG2-mic PCR product. (B) PCR screening of 20 colonies resulting from the CRISPR experiment. A 269 bp PCR product from the *JEN1* locus obtained with the primers DonJEN1_f / CheckJEN_r confirms a positive deletion. A negative (N) control PCR of a WT DNA with the same primers generates a fragment of 322 bp. (C) A 231 bp PCR product from the *DLD1* locus obtained with the primers CheckDLD1_f1 / DonDLD1_r confirms a positive deletion. A negative (N) control PCR of a WT DNA with the same primers generates a fragment of 285 bp. (D) A 327 bp PCR product from the *ROM2* locus obtained with the primers ROM2P3for / ROM2P3rev confirms a positive deletion. A negative (N) deletion control PCR of a WT DNA with the same primers generates a fragment of 378 bp.

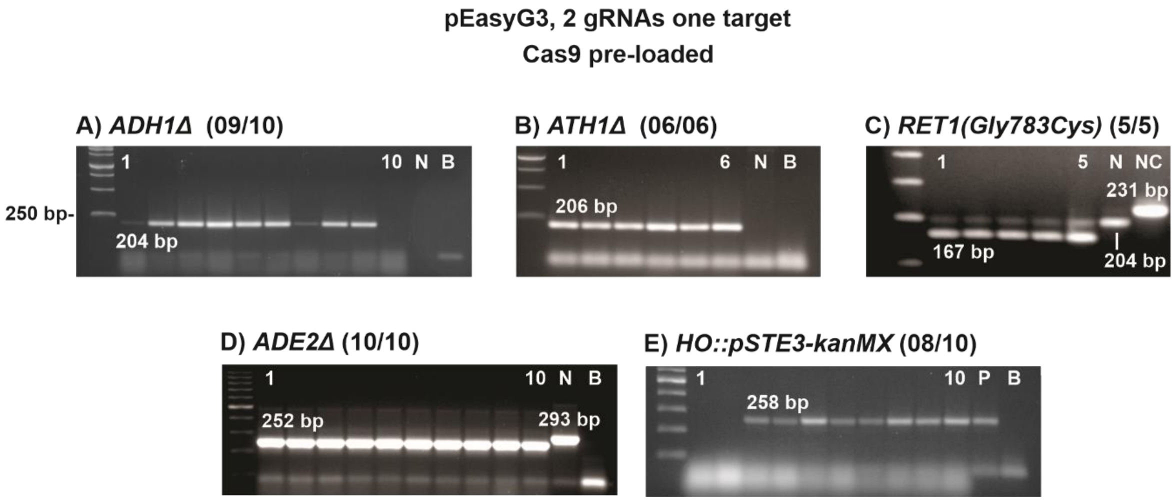


**Figure S14.** pEasyG3, CRISPR experiment using 2 gRNAs for a single deletion performed into Cas9 pre-loaded cells. 2% agarose gel electrophoreses of PCR fragments are shown from the screening of colonies for positive deletions. PCR blank controls (B) were included in the agarose gels, as well as negative (N) controls consisting of a PCR with the WT DNA (not deleted) and the same primers used to screen the transformants. (A) A 204 bp PCR product from *ADH1* obtained with primers CheckADH1_f / CheckADH1_r confirms deletions in 09 out of 10 colonies. Promega 1 Kb ladder was used as a molecular weight standard. (B) A 206 bp PCR product from *ATH1* obtained with primers ATHP2for / ATH1Er confirms deletions in 06 out of 06 colonies. Promega 1 Kb ladder was used as a molecular weight standard. (C) Point mutation introduced via CRISPR/Cas9 into *RET1* changes a glycine to a cysteine at the product position 783. A SalI restriction site was generated upon the CRISPR/Cas9 edit. Cleavage of a 231 bp PCR fragment (primers RET1SalI_f / RET1SalI_r) with SalI generates a 167 bp fragment only in the CRISPR modified DNA, while a 204 bp fragment is expected for the WT DNA (N) and for partially digested edited DNAs. NC, non-cleavage PCR product. (D) A 252 bp PCR product from the *ADE2* locus obtained with the primers ADE2f2 / ADE2r1 confirms deletions in 10 out of 10 colonies. A 293 bp negative (N) control corresponds to a PCR amplification of the WT locus. (E) CRISPR/Cas9-mediated insertion of the construct *pSTE3-KanMX* into the *HO* locus. The *pSTE3* and the *KanMX* PCR fragments (donors) were co-transformed with pEasyG3-hph and pEasyG3-mic PCR products. The fusion between the two fragments occurred in vivo upon *HO* insertion. A 258 bp PCR product obtained with the primers STE3-f2 / KAN5-rev confirms a fusion between the fragments. P, a positive PCR control with a strain in which the fusion has been already established.

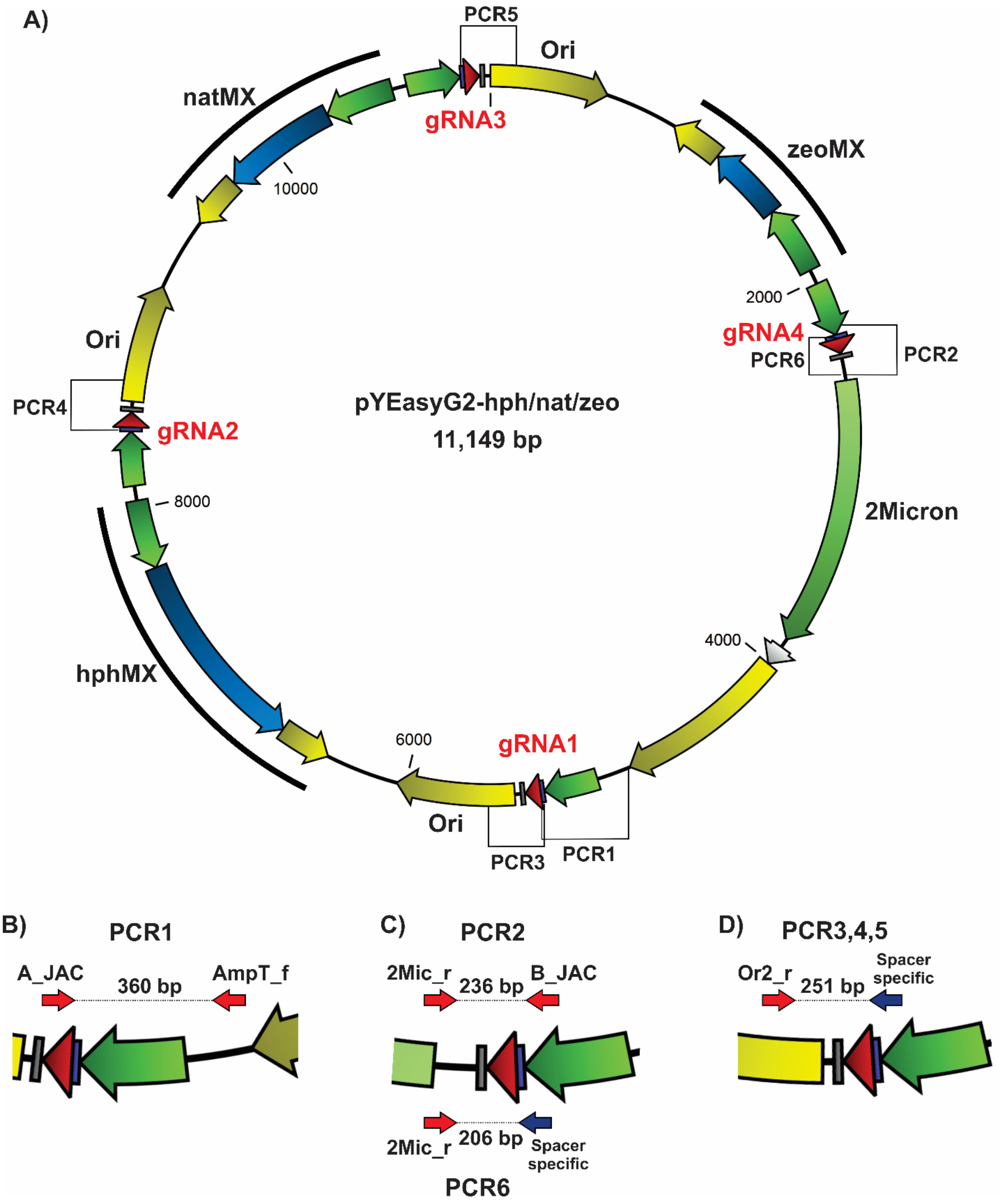


**Figure S15.** Map of the pYEasyG2 with four gRNAs assembled in vivo. (A) A structural map of the assembled plasmid is shown. PCRs for testing the assembly are indicated. (B-C) A detailed view is provided for the six PCRs with respective primers and product sizes. Spacer-specific primers (violet arrows) are 20-nts oligos matching the spacer sequence that serve to probe the exact assembly of a planned gRNA.

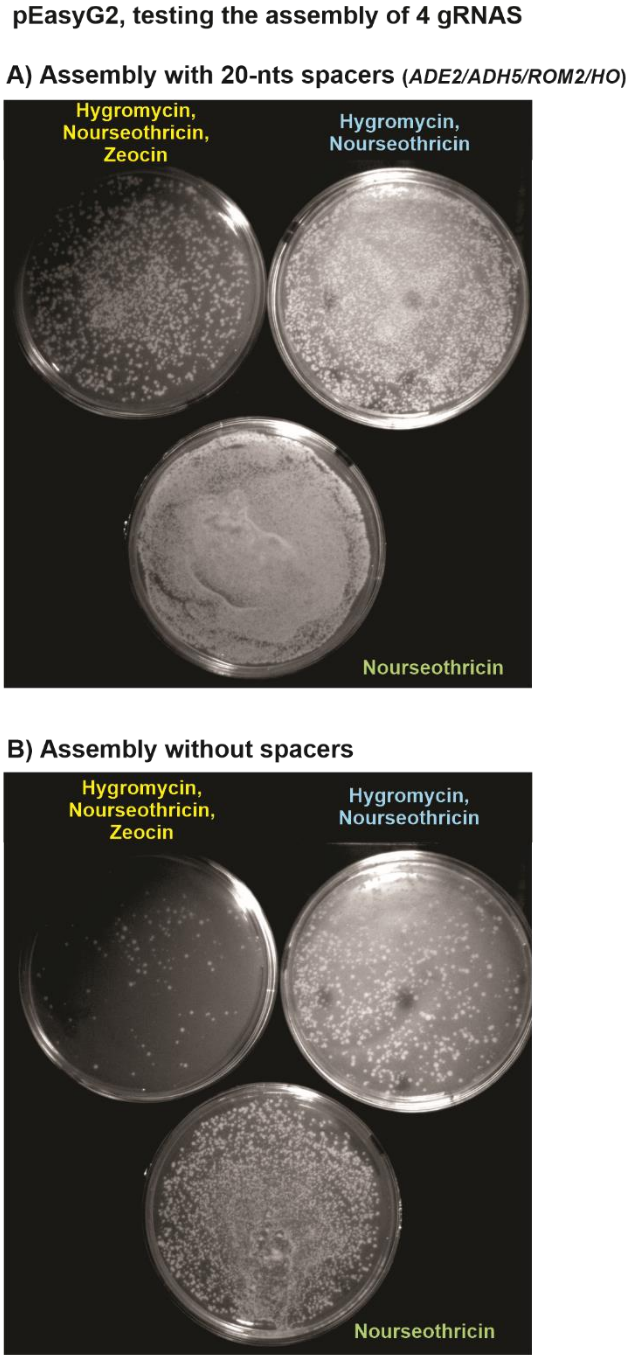


**Figure S16.** pEasyG2; antibiotic selection and spacers are needed for proper assembly of 4 gRNAs. (A) PCR fragments derived from pEasyG2-hph (gA_ADE2 / gB_ADH5), pEasyG2-nat (gA_ADH5 / gB_ROM2). pEasyG2-zeo (gA_ROM2 / gB_HO) and pEasyG2-mic (gA_HO / gB_ADE2) were co-transformed in equivolumetric amounts (15 µL from each PCR reaction) and plated onto solid medium with three, two, or one antibiotic, as indicated. Omitting antibiotic selection increases transformation rates likely due to gRNA misassemblies. (B) PCR fragments derived from pEasyG2-hph, pEasyG2-nat, pEasyG2-zeo, pEasyG2-mic (primers gA and gB without spacers) were co-transformed in equivolumetric amounts (15 µL from each PCR reaction) and plated onto solid medium with respectively three, two, or one antibiotic, as indicated. Transformation rates were much lower than observed for amplicons with spacers (A).

**pEasyG2, assembly of 4 gRNAs**

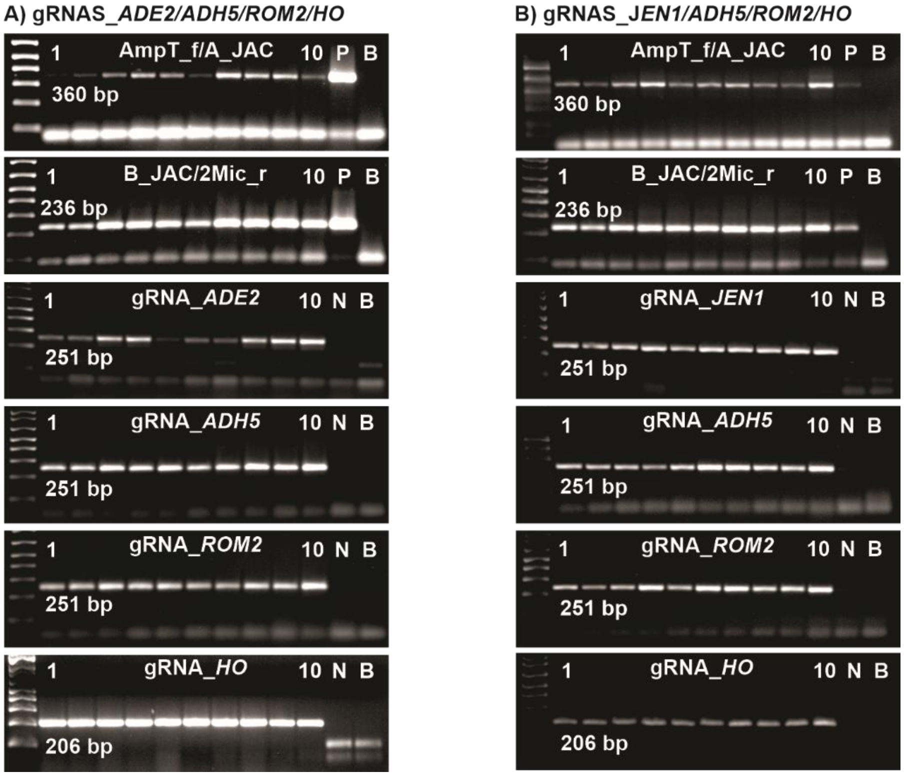


**Figure S17.** pEasyG2, testing the assembly of 4 gRNAs. The six PCRs shown in Figure S15 were used to probe the in vivo assembly of two pYEasyG2 replicons. 2% agarose gel electrophoreses of PCR fragments from 10 screened colonies are shown for each assemblage to confirm PCR amplifications. PCR blank controls (B) were included in the agarose gels. A 100 bp ladder was used as a molecular weight standard. Negative PCR controls (N) were conducted with spacer-specific primers used to amplify an unrelated pYEasyG2 having a different spacer. Positive PCR controls (P) amplify a common structure of an unrelated pYEasyG2. (A) Testing the assembly of gRNAs_*ADE2*/*ADH5*/*ROM2*/*HO* from colonies selected from the plate with three antibiotics (Figure S16, on the top, left). Amplification with spacer-specific primers (ADE2test, ADH5test, and ROM2test in combination with Or2_r, 251 bp; HOtest with 2Mic_r, 206 bp), confirm 100% assembly efficiency. (B) Testing the assembly of gRNAS_*JEN1*/*ADH5*/*ROM2*/*HO*. Amplification with spacer-specific primers (JEN1test, ADH5test, and ROM2test in combination with Or2_r, 251 bp; HOtest with 2Mic_r, 206 bp) confirm 100% assembly efficiency.

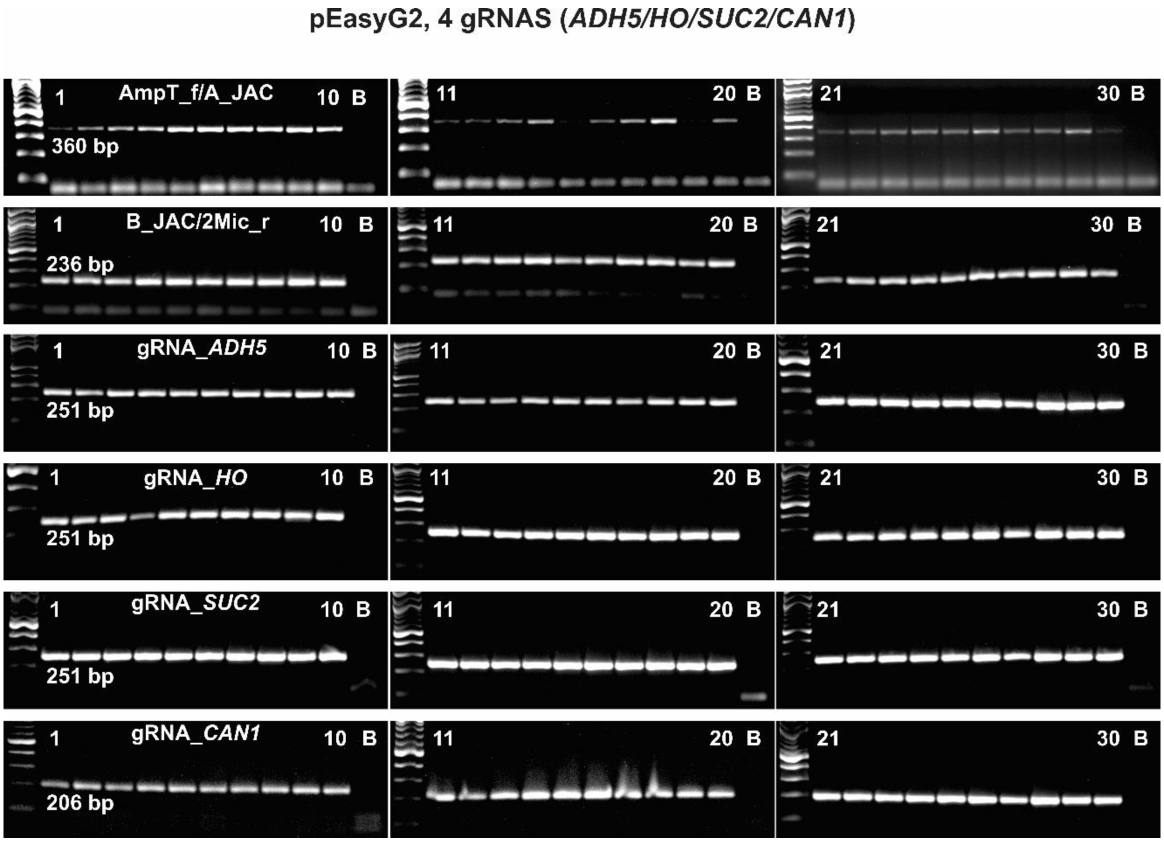


**Figure S18.** pEasyG2, testing the assembly of 4 gRNAs_*ADH5*/*HO*/*SUC2*/*CAN1*. The six PCRs shown on Figure S15 were used to probe the in vivo assembly of pYEasyG2_ *ADH5*/*HO*/*SUC2*/*CAN1*. 2% agarose gel electrophoreses of PCR fragments from 30 screened colonies (transformation triplicates) are shown to confirm PCR amplifications. PCR blank controls (B) were included in the agarose gels. A 100 bp ladder was used as a molecular weight standard. Amplification with spacer specific primers (ADH5test, HOtest, and SUC2test in combination with Or2_r, 251 bp; CAN1test with 2Mic_r, 206 bp) confirm 100% assembly in all 30 colonies tested.

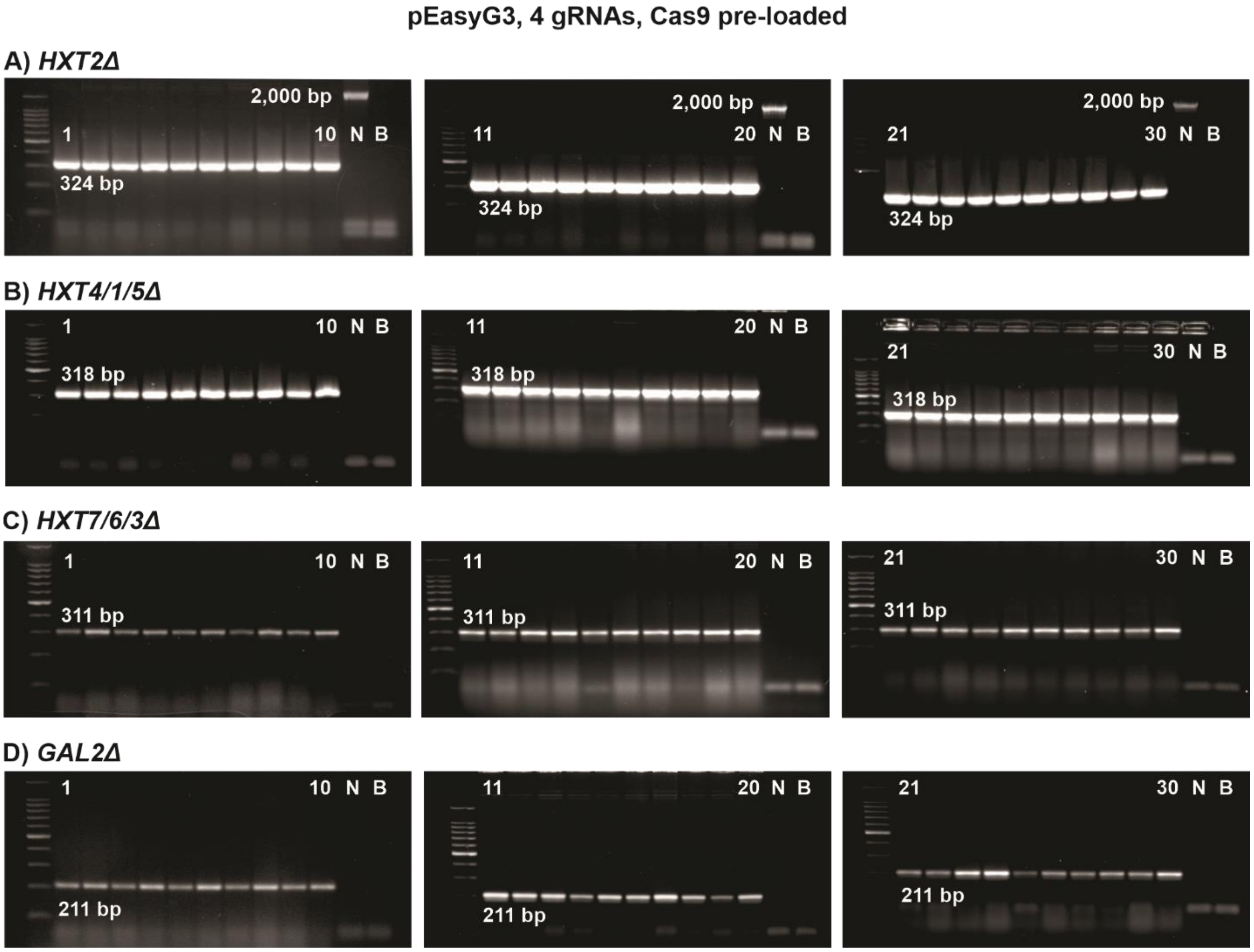


**Figure S19.** pEasyG3, 4 gRNAs for deletions of hexose transporters *HXT2*, *HXT4/1/5*, *HXT7/6/3*, and *GAL2*. 2% agarose gel electrophoreses of PCR fragments from 30 screened colonies are shown to confirm the targeted deletions. A negative PCR control (N) of the deletion, corresponding to a PCR product of the wild-type locus, and a PCR blank control (B) were included in the agarose gels. A 100 bp ladder was used as a molecular weight standard. (A) A 324 bp PCR product from the *HXT2* locus obtained with the primers HXT2pro_f / HXT2ter_r confirms a positive deletion. A negative deletion control PCR with primers HXT2pro_f / HXT2ter_r generates a fragment of 2,000 bp. (B) A 318 bp PCR product from the *HXT4*/*HXT1*/*HXT5* locus obtained with the primers HXT4pro_f / HXT5ter_r confirms a positive deletion. C) A 311 bp PCR product from the *HXT3*/*HXT6*/*HXT7* locus obtained with the primers HXT7pro_f / HXT3ter_r corresponds to a positive deletion. (D) A 211 bp PCR product from the *GAL2* locus with the primers GAL2pro_f / GAL2ter_r confirms a positive deletion.

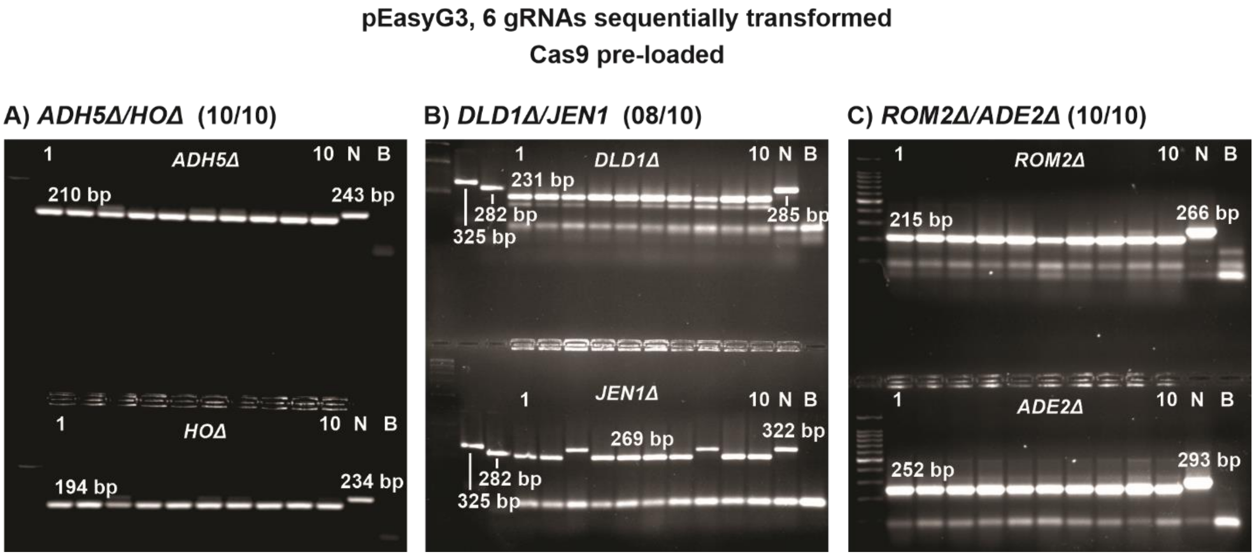


**Figure S20.** pEasyG3, sequential deletion of 6 targets. Screening of colonies obtained in each one of the three sequential rounds of transformation with pEasyG3-hph/nat/zeo and pEasyG3 fragments (see Figure 2D, main text). 2% agarose gel electrophoreses of PCR fragments from 10 screened colonies are shown to confirm the targeted deletions. A negative PCR control (N) of the deletion, corresponding to a PCR product of the wild-type locus, and a PCR blank control (B) were included in the agarose gels. A 100 bp ladder (A, C) and a 1 kb ladder (B) were used as molecular weight standards. (A) First round of transformation generating the *ADH5Δ*/*HOΔ* deletions. A 210 bp PCR product from *ADH5* (primers ADH5f2 / ADH5r2) and a 194 bp PCR product from *HO* (primers CheckHO_f / CheckHO_r) confirm the deletions. (B) Second round of transformation generating the *DLD1Δ*/*JEN1Δ* deletions. A 231 bp PCR product from the *DLD1* locus (primers CheckDLD1_f1 / DonDLD1_r) and a 269 bp PCR product from the *JEN1* locus (primers DonJEN1_f / CheckJEN_r) confirm the positive deletions. The two lanes next to the ladder are unrelated PCRs products of 325 bp and 282 bp, respectively, serving as molecular weight markers. (C) Third round of transformation generating the *ROM2Δ*/*ADE2Δ* deletions. A 215 bp PCR product from the *ROM2* locus (primers ROM2P3for / ROM2P3_r1) and a 252 bp PCR product from *ADE2* (primers ADE2f2 / ADE2r1) confirm the positive deletions.
